# Genetic diversity of endangered *Acropora spp.* corals in Bonaire: implications for restoration and gene banking

**DOI:** 10.64898/2026.10.27.762239

**Authors:** Allison Blanchette, Francesca Virdis, Sanne H. Tuijten, Pearl R. Rivers Key, Don R. Levitan, Sarah E. Lester, Andrew Rassweiler

**Affiliations:** Smithsonian Environmental Research Center, Edgewater, Maryland, USA; Department of Biological Science, Florida State University, Tallahassee, Florida, USA; Reef Renewal Foundation Bonaire, Kralendijk, Bonaire; Marine Biodiversity Group, Naturalis Biodiversity Center, Leiden, Netherlands

## Abstract

Species restoration is a valuable tool to combat biodiversity loss. However, maintaining genetic diversity in species restoration is challenging, particularly for species that can reproduce asexually, such as plants and corals, because a seemingly diverse collection of individuals could be composed of many clones of a few genotypes. Caribbean elkhorn and staghorn corals (*Acropora palmata, A. cervicornis*) are critically endangered reef-building corals that are a major target for restoration. To better understand the genetic diversity of an *Acropora spp.* restoration program and interpret local population genetic structure, we conducted microsatellite genotyping on the gene banks of 35 *A. palmata* lineages and 25 *A. cervicornis* lineages maintained by Reef Renewal Foundation Bonaire (RRFB) in the Caribbean Netherlands. We found that RRFB had high genotypic richness for *A. palmata* (0.971) and *A. cervicornis* (1.0) and estimated that they captured 88.9% and 74.7% of Bonaire’s microsatellite asymptotic allelic diversity, respectively. Compared to populations of *Acropora spp.* studied in other locations across the Caribbean, *A. palmata* in Bonaire had relatively low expected heterozygosity, while *A. cervicornis* was greater than approximately half of other populations. Across Bonaire and for both coral species, we did not find evidence for genetic structuring among spatially separated subpopulations, indicating that both species are likely well**-**connected around the island. Together, these results demonstrate the effectiveness of RRFB’s strategies to capture high genetic diversity and can be used to inform future initiatives on coral sexual breeding and assisted gene flow beyond Bonaire.

## Introduction

Globally, species are experiencing declines in population sizes, which can lead to losses of genetic diversity (Exposito-Alonso et al., 2022) through mechanisms such as genetic drift, reduced gene flow, and inbreeding. One tool to combat biodiversity loss is species restoration (i.e., supplementing or reintroducing a species through translocation, gardening, or breeding). However, genetic diversity can inadvertently be reduced in restoration by including a relatively small subset of the population’s genotypes and, intentionally or unintentionally, raising genotypes with high fitness in nursery settings (Holl, Luong & Brancalion, 2022). While protecting genetic diversity has been a primary focus in the conservation of species with severely depleted populations and inbreeding depression, such as the Florida panther (Johnson et al., 2010) and greater prairie-chicken (Capel et al., 2022), there has recently been increased attention to prioritize genetic diversity in restoration more broadly (Holl, Luong & Brancalion, 2022; Wei et al., 2023; O’Brien et al., 2025).

The growing emphasis on genetic diversity extends to coral restoration, with goals to conserve genetic diversity, create new genotypes through breeding, and strategically raise and outplant genotypes to support resilience to climate change (Vardi et al., 2021; Baums et al., 2022; Baker et al., 2025). There is a particular risk of unknowingly losing genetic diversity in corals because they can reproduce asexually (via fragmentation) as well as sexually. Thus, it is possible for a coral population to have relatively high abundance but low genetic diversity. For example, of 80 sampled individuals from the *Acropora cervicornis* population in Guadeloupe, all were asexual clones representing one single genotype (Japaud et al., 2015). Characterizing the genotypic diversity among individuals in a restoration program is key to understanding if trait variation is driven by genetic variation, as opposed to plasticity, and therefore the predictability of genotype performance in restoration. Investigating the genetic diversity of local and restored populations also informs restoration practitioners about the populations’ abilities to evolve and adapt to stressors such as climate change and disease outbreaks. Therefore, understanding the underlying genetic diversity of the restoration program is key to maximizing restoration success.

*Acropora spp.* corals are important foundation species that play an outsized role in providing habitat and food to reef fishes, relative to other hard coral species (Coker, Wilson & Pratchett, 2014). Caribbean *Acroporids* (*A. palmata, A. cervicornis,* Lamarck, 1816) have experienced large-scale declines due to stressors such as disease, bleaching, and local human impacts (Cramer et al., 2020), and are a major target in restoration (Young, Schopmeyer & Lirman, 2012). In the Florida Keys, there has been evidence of declines in genetic diversity of *A. palmata* in recent decades, prompting an urgent call to preserve the remaining genetic diversity of this species in nurseries (Williams, Miller & Baums, 2014; Williams et al., 2024). It has been generally suggested that 7-35 genotypes of *A. palmata* capture 50-90% of the wild (non-restored) population’s estimated asymptotic genetic diversity (Shearer, Porto & Zubillaga, 2009).

However, a wide range of genotypic richness values (proportion of sampled individuals that are unique genotypes) have been documented across wild *Acropora spp.* populations in the Caribbean, ranging from 0.01 to 1.0 (Baums, Miller & Hellberg, 2006; Williams, Miller & Baums, 2014; Porto-Hannes et al., 2015; Japaud et al., 2015; Canty et al., 2021). This variation in genotypic richness throughout the Caribbean can make it unclear for restoration programs how much genotypic diversity they will capture with local sampling, though practitioners do not always have the capacity to conduct lab-based genotyping to address this challenge.

*Acropora spp.* populations have previously been considered well-connected at large spatial scales spanning up to half of the Caribbean (Baums, Miller & Hellberg, 2005; Drury et al., 2016). However, there has also been evidence of more restricted gene flow (Vollmer & Palumbi, 2007) and localized population structuring observed in *A. cervicornis* (Garcia Reyes & Schizas, 2010; Drury et al., 2017; García-Urueña, Kitchen & Schizas, 2022). A better understanding of small-scale population genetic structure can help inform conservation by delineating management units and resolving how they map on governmental jurisdictions. It can also inform restoration outplanting and breeding by revealing if localized adaptation or inbreeding has occurred or is likely to occur.

In the southern Caribbean, the shallow coral reefs of the island of Bonaire (Caribbean Netherlands) were historically dominated by *Acropora spp.* (van Duyl, 1985). This dominance ended with Caribbean-wide declines of *Acropora spp.* in the 1980s, largely driven by disease (Aronson & Precht, 2001; Bruckner, 2003; Jackson et al., 2014; Boström-Einarsson et al., 2020; Cramer et al., 2020). Today, sparse stands of these critically endangered corals persist in Bonaire (Sommer et al., 2011; Crabbe et al., 2022; Miller et al., 2022; Kemenes van Uden, Meesters & Dogruer, 2023; Rolloos, Dogruer & Meesters, 2023), while they have relatively lower abundances in other parts of their range and have become functionally extinct in Florida (Manzello et al., 2025). A Marine Protected Area (MPA) was established in Bonaire in 1979 and surrounds the entire island from 0-60 meters depth and is managed by Stichting Nationale Parken Bonaire (STINAPA). The coral restoration organization Reef Renewal Foundation Bonaire (RRFB) has been propagating and outplanting lineages (assumed different genotypes) of *A. palmata* and *A. cervicornis* for over ten years (since 2012) with relatively high success (RRFB, 2024). Since the large-scale declines of *Acropora spp.*, only one site in Bonaire (Tailor Made) has been surveyed for genetic diversity of *A. palmata* (Baums, Miller & Hellberg, 2005, 2006), a site which demonstrated relatively high genotypic richness (32 genets out of 43 individuals) compared to other localities throughout the Caribbean.

In this study we conducted microsatellite genotyping of 35 lineages of *A. palmata* and 25 lineages of *A. cervicornis* collected in Bonaire by RRFB for restoration. Our primary objectives were to characterize the genetic diversity of *Acropora spp.* in RRFB’s gene bank and evaluate how much of Bonaire’s genetic diversity has been captured by RRFB. We additionally used this data to interpret genetic structuring among potential subpopulations on Bonaire, based on allelic fixation and differentiation, inbreeding, and relatedness. Despite the utility of sequencing approaches, we chose to use microsatellites because they are cost-effective and function comparably to Single Nucleotide Polymorphisms (SNPs) in determining genetic diversity and structuring among populations (Lemopoulos et al., 2019; Sunde et al., 2020; Zimmerman, Aldridge & Oyler-McCance, 2020). Microsatellites have also been used to genotype lineages in multiple Caribbean *Acropora spp.* restoration programs (Carne, Kaufman & Scavo, 2016; O’Donnell et al., 2018; Miller et al., 2019; Calle-Triviño et al., 2020), making them more relevant for comparison to other restoration efforts.

## Materials & Methods

### Lineage collection

Reef Renewal Foundation Bonaire (RRFB) has collected fragments of *A. palmata* and *A. cervicornis* from wild coral colonies (i.e., donor colonies) in Bonaire from 2012 to present (Nature permit issued by Openbaar Lichaam Bonaire, archive number R&O2023000162). RRFB propagates the collected coral fragments asexually in underwater nurseries, keeping track of descendants (hereafter lineages), as a means of gene banking and raising lineages for outplanting. Lineages’ donor colonies were located across the full span of Bonaire’s leeward western coastline and in Lac Bay on the windward east coast (Fig. 1a). We categorized donor colony locations into five potential subpopulations defined by their separation by both physical distance and geographic features: North leeward, Central leeward, South leeward, Klein Bonaire, and Lac Bay. Klein Bonaire is a small uninhabited island approximately 750 meters from Bonaire’s leeward west coast, separated from the coast by a deep (>140m) marine channel (Becking & Meesters, 2014; De Meyer et al., 2022). Lac Bay is a sheltered shallow bay on Bonaire’s windward east coast. The potential subpopulations on mainland Bonaire were all separated from each other by at least 2.5 kilometers and separated from Klein Bonaire by at least one kilometer. Geographic features also separated most of the potential subpopulations, including the isolation of Lac Bay, the marine channel separating Klein Bonaire from mainland Bonaire, and the point between central and south Bonaire (Punt Vierkant). Aside from Lac Bay, collections were not conducted on Bonaire’s windward east coast due to difficult conditions (i.e., strong wave action) and the lower coral cover as a result of high physical disturbance (van Duyl, 1985; Sommer et al., 2011). Each lineage RRFB collected was assigned to a subpopulation according to the location of its donor colony, though it is possible that a lineage had asexual clones present in other locations due to fragmentation and transportation by strong wave action. For *A. palmata* (N = 35 lineages), the subpopulation sample sizes were: North (n = 6), Central (n = 8), South (n = 8), Klein (n = 4), and Lac Bay (n = 9). For *A. cervicornis* (N = 25 lineages), the subpopulation sample sizes were: North (n = 7), Central (n = 2), South (n = 11), Klein (n = 3), Lac Bay (n = 1), and Unknown (n = 1). The *A. cervicornis* lineage with unknown donor colony location was retained in all analyses that did not require an assigned subpopulation but was excluded from analyses that required a subpopulation.

**Figure 1.**
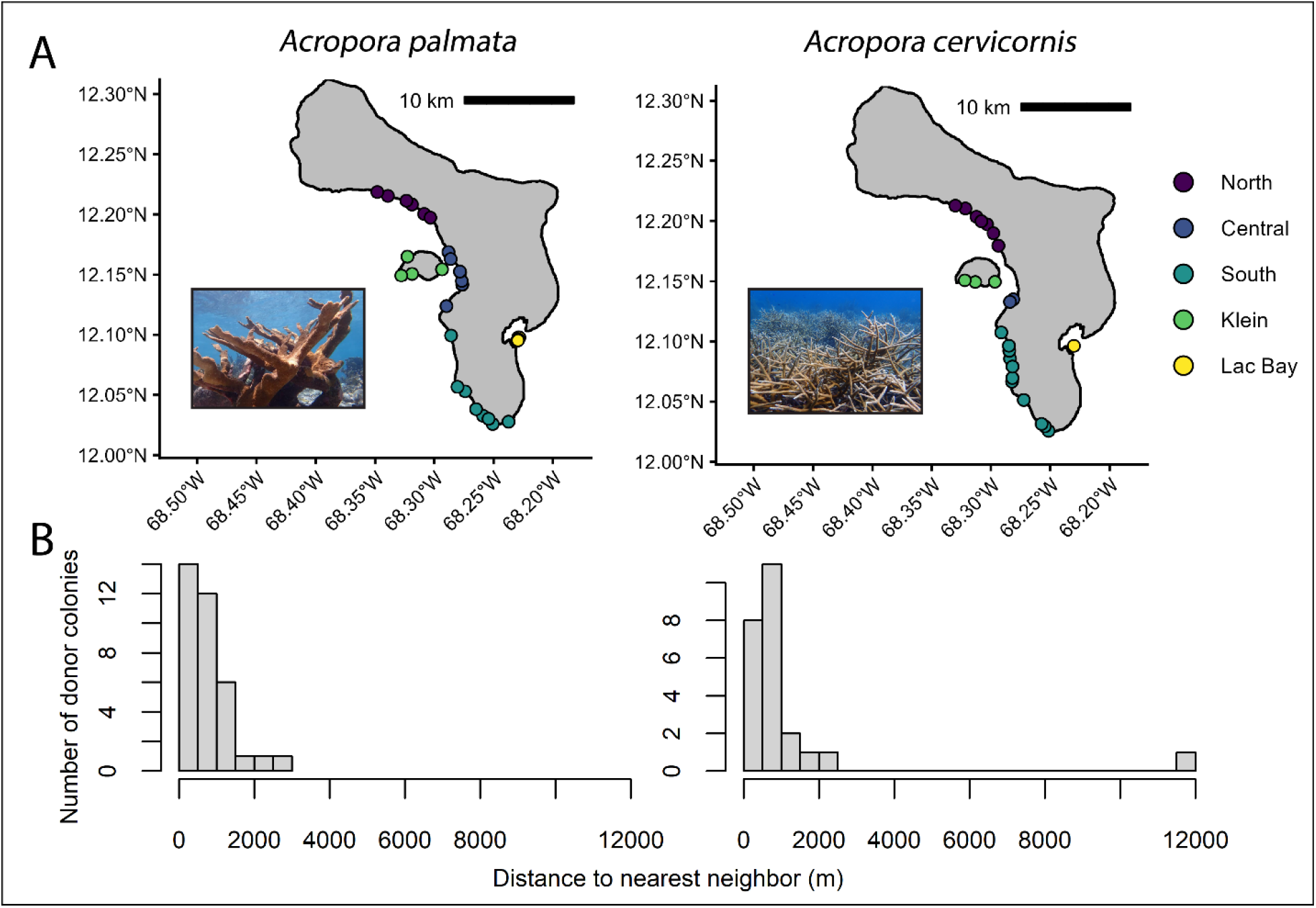
Maps of lineage donor-colony locations and histograms of nearest-neighbor distances among donor-colonies. (A) Maps of lineage donor-colony locations on Bonaire for *Acropora palmata* and *A. cervicornis*. Points are color-coded based on assigned sub-population. For *A. palmata* (N = 35 lineages), the sub-population sample sizes were: North (n = 6), Central (n = 8), South (n = 8), Klein (n = 4), and Lac Bay (n = 9). For *A. cervicornis* (N = 25 lineages), the sub-population sample sizes were: North (n = 7), Central (n = 2), South (n = 11), Klein (n = 3), and Lac Bay (n = 1), and Unknown (n = 1). Map shapefile sourced from Humanitarian Data Exchange (HDX, 2017). (B) Histograms depict distribution of distances of nearest neighbors among donor-colonies. The nearest neighbor of the single *A. cervicornis* donor-colony in Lac Bay was on the southern tip of Bonaire, approximately 11,800m away.

All chosen donor colonies were spatially separated from one another to reduce the probability of collecting genetically identical clones (Fig. 1b). Sixty-three percent (22 out of 35) of the *A. palmata* lineages were collected from donor colonies >300 meters away from the next nearest *A. palmata* donor colony. However, the other 13 lineages were collected from donor colonies 3-90 meters away from other donor colonies, primarily in Lac Bay because of the spatial clustering of its limited stands of *A. palmata*. All *A. cervicornis* donor colonies were >300 meters away from other *A. cervicornis* donor colonies. Lineages were assumed to be genetically distinct genotypes, but this had not been confirmed until the present study.

### Genetic sampling

To conduct the genetic analyses, we collected coral tissue samples from RRFB’s full gene bank of *A. palmata* and *A. cervicornis* in 2022 and 2024. This research was permitted by Openbaar Lichaam Bonaire (Nature permit, archive number R&O2023000162) and coral tissue sample collection and export from Bonaire were permitted by the Convention on International Trade in Endangered Species (CITES, permit numbers BQ/2022/002, BQ/2024/003). In total, we sampled 35 *A. palmata* lineages and 25 *A. cervicornis* lineages. For each lineage, we collected one 1cm^2^ clipping of the apical tip from two separate fragments in a nursery tree (each tree maintained one lineage). The samples were immediately rinsed with filtered freshwater (e.g., bottled or tap) and placed in preservative. In 2022, samples were preserved in salt-saturated DMSO buffer (Gaither et al., 2011): 30 *A. palmata* lineages, 24 *A. cervicornis* lineages. In 2024, we sampled six additional newly collected lineages (five new *A. palmata* lineages, one new *A. cervicornis* lineage). Preserved samples were kept in the dark at -29°C until DNA extraction.

Based on relatively low DNA yields from *A. cervicornis* in the initial 2022 sampling (median 0.9μg/mL), we also tested the effectiveness of DNA/RNA Shield (Zymo Research) as a preservative in comparison to DMSO in 2024. Across a subset of five *A. cervicornis* samples that had replicates stored in both preservative types, the DNA yields were nearly an order of magnitude greater when preserved in DNA Shield (median 20.7μg/mL) than DMSO (median 3.6μg/mL) (Supplemental Material S1, Table S1).

### Microsatellite genotyping

To extract the samples’ DNA, we used the Qiagen DNeasy tissue kit, following modified methods for *Acropora spp.* (Baums & Kitchen, 2020). We ran multiplex PCRs with 11 microsatellite loci developed and published by Baums, Hughes & Hellberg (2005) and Baums et al. (2009), split across three separate mixes (**Mix A**: 182, 0585, 1195, **Mix B**: 5047, 207, 181, 6212, **Mix C**: 0513, 2637, 1490, 166). To differentiate among loci in the multiplex, we used fluorescently labeled universal primer tails (Blacket et al., 2012). Primer sequences are available in Supplemental Material S2, Table S2. We followed the Qiagen Multiplex Mix thermocycler settings, with annealing temperatures at 54°C for Mixes A and B, and 53°C for Mix C, for 35 cycles. We processed the PCR product through fragment analysis with GeneScan 500 LIZ size standard (Applied Biosystems) at the Florida State University DNA Sequencing Facility using a 3730 DNA analyzer (Applied Biosystems). To account for possible error in the PCR and fragment analysis steps, we ran three replicates of each extracted DNA sample per lineage. Alleles were scored using Geneious Prime 2025.0.3 (https://www.geneious.com).

### Genetic diversity in gene bank

We conducted all analyses in R (4.4.3) using R Studio (2026.01.0). To convert the allele table data formatted as comma-separated values (csv) into a genind format compatible with relevant R packages, we used the function *df2genind()* from adegenet (Jombart, 2008). To determine whether each lineage was a unique genotype or a clone of another lineage, we manually assessed allele identities across all 11 loci. If a lineage’s combination of allele values across all loci was unique from all other lineages, then we determined it to be a unique Multi-Locus Genotype (MLG). We evaluated confidence in MLG identification with three methods, by: 1.) determining the number of loci necessary to identify an MLG (*A. palmata*: four loci, *A. cervicornis*: five loci), 2.) estimating the probability of two lineages sharing an identical MLG via sexual reproduction rather than asexual fragmentation (*A. palmata* P_sex_ <4^-5^, *A. cervicornis* P_sex_ <2^-3^ for all lineages except G01 which was ultimately excluded due to triploidy), and 3.) determining if any MLGs differed solely due to somatic mutation, using a Bruvo’s genetic distance threshold of 0.1 (Bruvo’s genetic distance > 0.1 for all lineages except two identical *A. palmata* clones). For further detail on analyses of confidence in MLG identification, see Supplemental Material S3 and Fig. S1. In all subsequent analyses for this study, only loci with <15% missing data were used (*A. palmata*: 11 loci, *A. cervicornis*: six loci).

To characterize genetic diversity in RRFB’s gene bank, we calculated genotypic richness (proportion of sampled lineages that were unique MLGs) and microsatellite allelic diversity (number of unique alleles, heterozygosity), both of which are necessary to measure as they capture different dimensions of genetic diversity. Genotypic richness measures number and frequency of genetically unique individuals (i.e., clonal redundancy in the gene bank), while microsatellite allelic diversity measures number and frequency of unique alleles across the different MLGs. Microsatellite allelic diversity can be used to estimate how much of the wild population’s allelic diversity is represented in the gene bank (i.e., asymptotic allelic diversity). We calculated number of unique alleles using poppr and the expected and observed heterozygosity using hierfstat (Goudet & Jombart, 2022) (Table 1). We conducted rarefactions with iNEXT (Chao et al., 2014; Hsieh, Ma & Chao, 2016) to estimate how much of Bonaire’s *Acropora spp.* genetic diversity has been captured by RRFB. While iNEXT is designed to rarefy species diversity as a function of sample size, we leveraged it to rarefy the total number of unique alleles across loci as a function of MLG sample size in order to estimate allelic asymptotic diversity (1,000 bootstrap iterations). To standardize our rarefaction results, we averaged the total number of unique alleles by the number of microsatellite loci assessed for each species (*A. palmata*: 11 loci, *A. cervicornis*: six loci). To check for consistency in rarefaction patterns across loci, we used the pegas package to conduct rarefactions on number of unique alleles for each microsatellite locus individually.

**Table 1.** Genetic diversity at 11 microsatellite loci for *Acropora palmata* and six microsatellite loci for *Acropora cervicornis.* N_a_: Number of unique alleles, H_e_: expected heterozygosity, H_o_: observed heterozygosity, p: significance of deviation from Hardy-Weinberg equilibrium (alpha = 0.05).

| **Species** | **Locus** | **N<sub>a</sub>** | **H<sub>o</sub>** | **H<sub>e</sub>** | **p** |
|---|---|---|---|---|---|
| *Acropora palmata* | 182 | 14 | 0.68 | 0.61 | 0.96 |
|  | 0585 | 7 | 0.59 | 0.67 | 0.83 |
|  | 1195 | 6 | 0.61 | 0.66 | 0.65 |
|  | 5047 | 6 | 0.71 | 0.73 | 0.63 |
|  | 207 | 13 | 0.76 | 0.87 | 0.02 |
|  | 181 | 3 | 0.21 | 0.29 | 0.05 |
|  | 6212 | 13 | 0.82 | 0.85 | 0.31 |
|  | 0513 | 7 | 0.29 | 0.34 | 0.21 |
|  | 2637 | 2 | 0.53 | 0.48 | 0.71 |
|  | 1490 | 7 | 0.62 | 0.71 | 0.25 |
|  | 166 | 11 | 0.59 | 0.71 | 0.001 |
|  | Average | 8.09 | 0.58 | 0.63 | NA |
| *Acropora cervicornis* | 182 | 8 | 0.82 | 0.72 | 0.83 |
|  | 0585 | 7 | 0.67 | 0.72 | 0.26 |
|  | 5047 | 5 | 0.71 | 0.76 | 0.69 |
|  | 181 | 6 | 0.75 | 0.84 | 0.53 |
|  | 1490 | 5 | 0.59 | 0.68 | 0.58 |
|  | 166 | 5 | 0.65 | 0.57 | 0.74 |
|  | Average | 6.00 | 0.70 | 0.72 | NA |

To understand how the genetic diversity in RRFB’s gene bank compared to other *Acropora spp.* populations, we compared our results to those reported in prior studies on Caribbean *Acropora spp.* population genetics (for literature search methods, see Supplemental Material S4). To do this, we averaged the number of unique alleles and expected heterozygosity across a subset of loci that had available data reported in all studies being compared, which included four loci in *A. palmata* (166, 181, 182, 207) and four loci in *A. cervicornis* (166, 181, 182, 0585). All data from prior studies were gathered from the main paper or supplemental section of peer-reviewed publications.

To assess if there were still more *Acropora spp.* genotypes in Bonaire yet to be sampled by RRFB, we compared the coordinates of RRFB’s collection locations to mapped *Acropora spp.* stands (i.e., individual colonies and dense thickets). The mapping surveys were conducted by Wageningen Marine Research and STINAPA in 2022, which surveyed 19 areas along the west coast and Klein Bonaire (146 hectares total shelf area) that covered approximately 13% of Bonaire’s coastline (Kemenes van Uden, Meesters & Dogruer, 2023; Rolloos, Dogruer & Meesters, 2023). Using mapping surveys, we calculated the percent of mapped stands within the surveyed areas that RRFB had not collected from and identified additional stands that RRFB collected from that were not identified in the mapping surveys.

### Population structure

To interpret population structuring within Bonaire based on genetic data from RRFB’s gene bank, we compared genetic diversity among the five hypothesized subpopulations: North, Central, South, Klein Bonaire, and Lac Bay. The sample size of MLGs per potential subpopulation was identical to the number of lineages per subpopulation described above in Lineage collection, except for two lineages that were excluded: one *A. palmata* lineage from Lac Bay because it shared an identical MLG identity with another lineage in Lac Bay, and *A. cervicornis* lineage G01 from Lac Bay because it had a relatively high occurrence of triploidy (three loci). *A. cervicornis* population structuring analyses thus excluded Lac Bay because lineage G01 was the only individual collected in Lac Bay. Additionally, sample sizes were small for *A. cervicornis* Central (n = 2) and Klein (n = 3), making the analyses of *A. cervicornis* low-powered and primarily exploratory. Overall, while RRFB’s lineage collection strategies were more targeted at maximizing genotypic richness than investigating population genetics (Meirmans, 2015), we were able to interpret if there was any strong population structuring from RRFB’s gene bank because of the relatively high number of MLGs it captured (Shearer, Porto & Zubillaga, 2009).

To visualize genetic differences among potential subpopulations, we ran a Principal Component Analysis (PCA) on allele frequencies with the R packages ade4 and adegenet (Chessel, Dufour & Thioulouse, 2004; Jombart, 2008), imputing missing data based on mean allele frequencies. To explore if there was any clustering within Bonaire that better represented the data than the five potential subpopulations that we defined, we used *find.clusters()* in adegenet (Jombart, 2008; Jombart, Devillard & Balloux, 2010) as well as Structure Selector (Li & Liu, 2018) at https://lmme.ac.cn/StructureSelector/ (full methods in Supplemental Material S5, Fig. S2-S4). To test the results from Structure Selector, we inspected STRUCTURE analysis outputs when the number of clusters (K) was set equal to the optimal K identified in the ΔK plots from Structure Selector.

To quantify genetic differences among potential subpopulations, we calculated pairwise G’’_ST_ among subpopulations to compare nearness to fixation, because G’’_ST_ accounts for high heterozygosity that is common in microsatellites and small sample size better than other F-statistics (Hedrick, 2005; Meirmans & Hedrick, 2011). We also calculated Jost’s D (Jost, 2008) as a measure of allelic differentiation. To test if G’’_ST_ and Jost’s D were significantly greater than 0, we obtained 95% confidence intervals for the global values via bootstrapping (Chao et al., 2008) with 1,000 iterations. We deemed G’’_ST_ and Jost’s D to be significantly greater than zero if the lower confidence interval was greater than zero. These analyses were conducted using mmod (Winter, 2012). We also calculated G_ST_ (Nei, 1973) in mmod and θ (Weir & Cockerham, 1984) in hierfstat, as they are other commonly-used F-statistics.

To understand how MLGs were related to each other within and across potential subpopulations, we calculated triadic maximum likelihood (TrioML) estimator of relatedness (Wang, 2007) because it has been found to outperform other estimators of relatedness on microsatellite data (Wang, 2007; Taylor, 2015), using the related package (Pew et al., 2015). To determine if observed relatedness values were greater than expected, which is indicative of inbreeding or non-random mating, we generated distributions of expected relatedness values via bootstrapping (1,000 iterations) for each potential subpopulation and for Bonaire overall. We calculated inbreeding coefficients (F_IS_) using hierfstat.

## Results

All 11 microsatellite loci were successfully genotyped across ≥85% of lineages (i.e., <15% of lineages had missing data) in *A. palmata* (182, 0585, 1195, 5047, 207, 181, 6212, 0513, 2637, 1490, 166) and at six loci in *A. cervicornis* (182, 0585, 5047, 181, 1490, 166). We found no evidence of peak call errors, linkage disequilibrium, or null alleles in the loci with <15% missing data (Supplemental Material S6). Three *A. cervicornis* lineages were triploid at locus 182 (lineages G13, G14, G25), which exhibited triploidy in Nylander-Asplin et al. (2021) as well. We excluded the apparent triploid alleles at locus 182 from analyses because of the dependence of most population genetic analyses on diploid data. *A. cervicornis* lineage G01 (Lac Bay subpopulation) also exhibited triploidy at three loci (181, 1490, and 6212), so we excluded this lineage altogether from analyses because of uncertainty in allele dosage at the other loci (Dufresne et al., 2014; Meirmans, Liu & van Tienderen, 2018).

### Genetic diversity in gene bank

We found that 34 of the 35 lineages of *A. palmata* in RRFB’s gene bank were unique Multi-Locus Genotypes (MLGs) and all 25 lineages of *A. cervicornis* were unique MLGs. Thus, there was a genotypic richness (proportion of sampled lineages that are unique MLGs) of 0.971 for *A. palmata* and 1.000 for *A. cervicornis*. The two *A. palmata* lineages that were identical MLGs were originally collected from donor colonies that were 6.8 meters apart from one another in Lac Bay and were likely asexual clones from the same original MLG. Interestingly, another pair of *A. palmata* lineages were collected within just three meters of each other in Lac Bay and were different MLGs, emphasizing the potential for genetic diversity even on small spatial scales.

There was a relatively high number of unique alleles and expected heterozygosity for most loci, and all loci except for 207 and 166 in *A. palmata* were in Hardy-Weinberg Equilibrium (Table 1). We retained loci 207 and 166 in *A. palmata* analyses because we estimated, using three independent tests, that null allele frequencies for these loci were not significantly greater than zero, except for locus 166 in the Micro-checker test (van Oosterhout et al., 2004). However, sensitivity testing showed no difference in results when locus 166 was excluded (Supplemental Material S6). To estimate how much of Bonaire’s genetic diversity of *Acropora spp.* was captured in RRFB’s gene bank, we used rarefaction to calculate the average number of unique alleles (i.e., allelic richness) as a function of the number of MLGs sampled (Fig. 2). The gene bank of *A. palmata* had 8.09 unique alleles per microsatellite locus on average (N = 11 loci) and we estimated that the asymptotic diversity was 9.10 unique alleles per locus.

**Figure 2.**
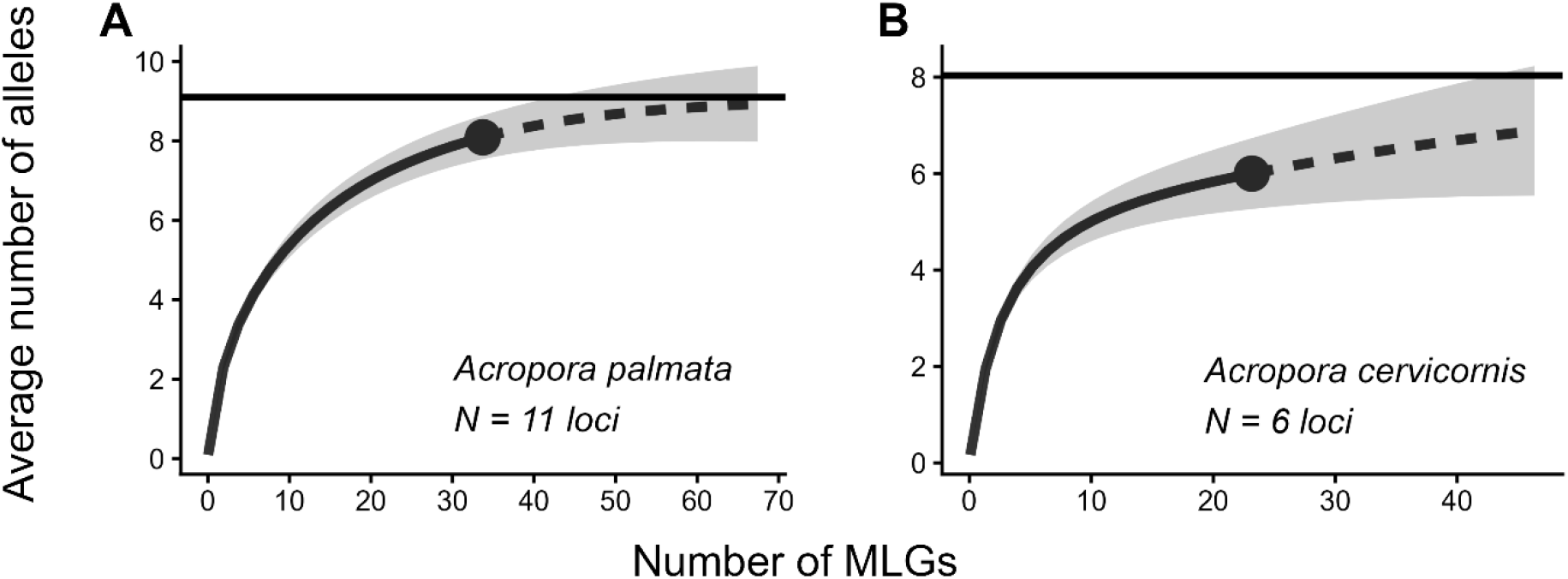
Rarefaction curves of average number of unique microsatellite alleles as a function of Multi-Locus Genotypes (MLGs) sampled. Number of unique alleles was averaged across 11 loci for *Acropora palmata* (A) and six loci for *Acropora cervicornis* (B). Solid rarefaction curve represents observed data while dotted curve represents extrapolated predictions, gray shading on rarefaction curve represents 95% confidence intervals, solid horizontal line represents predicted asymptotic allelic richness.

Thus, RRFB captured 88.9% of the asymptotic microsatellite allelic diversity for *A. palmata*. The gene bank of *A. cervicornis* had 6.00 unique alleles per microsatellite locus on average (N = 6 loci) and we estimated that the asymptotic diversity was 8.03 unique alleles per locus. Thus, RRFB captured 74.7% of the asymptotic microsatellite allelic diversity for *A. cervicornis*. In addition to average allelic richness across loci, we also rarified number of unique alleles *per locus.* The majority of loci exhibited similar patterns of approaching asymptotic diversity (Supplemental Material S7, Fig. S5 and S6).

The expected heterozygosity of *A. palmata* in RRFB’s gene bank was lower than the expected heterozygosity of all other previously studied *A. palmata* populations (lowest of 15 total populations) that were included in this comparison (Table 2). The expected heterozygosity of *A. cervicornis* was greater than 43% (three out of eight total populations) of other previously surveyed populations, including Puerto Rico, Navassa, and U.S. Virgin Islands. Both *A. palmata* and *A. cervicornis* had a greater average number of unique alleles than >20% of other populations, though number of unique alleles was also positively associated with MLG sample size. Interestingly, the expected heterozygosity and number of unique alleles of *A. palmata* were closely comparable to a previous study that surveyed *A. palmata* at one site in Bonaire (Tailer Made) in 2005 (Baums, Miller & Hellberg, 2005).

**Table 2.** Comparisons of genetic diversity between the present study and prior studies on *Acropora spp.* population genetics. N_a_: average number of unique alleles across loci, N_g_: number of MLGs, H_e_: average expected heterozygosity across loci. Averages were taken across a subset of loci that had available data reported in all studies being compared: four loci in *A. palmata* (166, 181, 182, 207) and four loci in *A. cervicornis* (166, 181, 182, 0585). Due to subsetting to four common loci, values in Table 2 are not directly comparable with Table 1 for the present study or with the prior studies if they used other microsatellite loci in addition to the common four. Table is sorted in ascending order of H_e_. Present study is bolded, other studies’ full citations in References list.

| **Population** | **N<sub>a</sub>** | **N<sub>g</sub>** | **H<sub>e</sub>** | **Reference** |
|---|---|---|---|---|
| *Acropora palmata* |  |  |  |  |
| **Bonaire** | **10.25** | **34** | **0.621** | **Present study** |
| Bonaire | 8.50 | 32 | 0.625 | Baums et al., 2005b |
| St Vincent and the Grenadines | 13.00 | 166 | 0.646 | Baums et al., 2005b |
| U.S. Virgin Islands | 11.75 | 41 | 0.677 | Baums et al., 2005b |
| Curacao | 15.00 | 100 | 0.700 | Baums et al., 2005b |
| Panama | 10.25 | 39 | 0.724 | Baums et al., 2005b |
| Puerto Rico | 14.75 | 90 | 0.739 | Baums et al., 2005b |
| Mona Island | 12.50 | 36 | 0.743 | Baums et al., 2005b |
| Florida Keys, 2006 surveys | 13.25 | 38 | 0.755 | Williams et al, 2014 |
| Florida Keys, 2010 surveys | 12.75 | 46 | 0.763 | Williams et al, 2014 |
| Guadeloupe | 8.25 | 20 | 0.775 | Japaud et al., 2015 |
| Florida Keys | 12.25 | 34 | 0.779 | Baums et al., 2005b |
| Mexico | 5.75 | 7 | 0.786 | Baums et al., 2005b |
| Navassa | 14.25 | 68 | 0.787 | Baums et al., 2005b |
| Bahamas | 16.25 | 96 | 0.789 | Baums et al., 2005b |
| *Acropora cervicornis* |  |  |  |  |
| Puerto Rico | 4.00 | 7 | 0.581 | Baums et al., 2010 |
| Navassa | 5.00 | 6 | 0.670 | Baums et al., 2010 |
| U.S. Virgin Islands | 8.25 | 27 | 0.683 | Baums et al., 2010 |
| **Bonaire** | **6.50** | **24** | **0.713** | **Present study** |
| Bahamas | 8.50 | 13 | 0.734 | Baums et al., 2010 |
| Honduras | 10.50 | 53 | 0.758 | Baums et al., 2010 |
| Florida Reef Tract | 11.00 | 172 | 0.772 | Baums et al., 2010 |
| Honduras | 11.25 | 161 | NA | Canty et al., 2021 |

While there was substantial *Acropora spp.* genetic diversity in RRFB’s gene bank, we also estimated what percent of remaining *Acropora spp.* stands (i.e., individual colonies and dense thickets) in Bonaire could be sources of new alleles, via new genotypes, for RRFB to more closely approach the asymptotic maximum diversity implied by analysis of their existing gene bank. Of the *Acropora spp.* mapped across 13% of Bonaire’s coastline (146 hectares total shelf area) in 2022 (Kemenes van Uden, Meesters & Dogruer, 2023; Rolloos, Dogruer & Meesters, 2023), RRFB had collected from 10.4% of the mapped *A. palmata* stands (five out of 48) (Supplemental Material S8, Fig. S7). Additionally, 28 *A. palmata* lineages were collected from donor colonies outside of surveyed areas and two lineages were from colonies that were within surveyed areas but were not mapped (e.g., colony died or was moved by wave action in between collection and mapping events). For *A. cervicornis*, RRFB had collected from 12.5% of the mapped stands (five out of 40). Fourteen *A. cervicornis* lineages were from donor colonies outside of surveyed areas and five were from unmapped stands within surveyed areas.

### Minimal population structuring

We did not detect any strong genetic structuring among potential subpopulations for either species (Fig. 3), based on measures of nearness to fixation, differentiation, and relatedness. Global comparison values of G’’_ST_ and Jost’s D were not significantly greater than zero for either species (Table 3). Pairwise comparisons of G’’_ST_ and Jost’s D between potential subpopulations were low in *A. palmata* (<0.03) and slightly higher in *A. cervicornis* (<0.20), though analyses of *A. cervicornis* were primarily exploratory and should be interpreted with caution due to low and unbalanced sample sizes of the potential subpopulations (e.g., Central = 2, Klein = 3). Pairwise G_ST_ and θ were also low in both species (<0.07, Supplemental Material S9, Table S3). In *A. palmata*, relatedness estimates (TrioML) were relatively low (<0.1) within and across the potential subpopulations, while relatedness estimates in *A. cervicornis* were slightly higher, with overall relatedness at 0.128 (Table 4). In both species, observed relatedness estimates were not greater than expected (via 1,000 bootstrapping iterations, Supplemental Material S10, Fig. S8 and S9) within and across potential subpopulations. In *A. palmata*, inbreeding coefficients (F_IS_) ranged from 0-0.168, while *A. cervicornis* values were slightly lower (F_IS_ ≤0.1).

**Figure 3.**
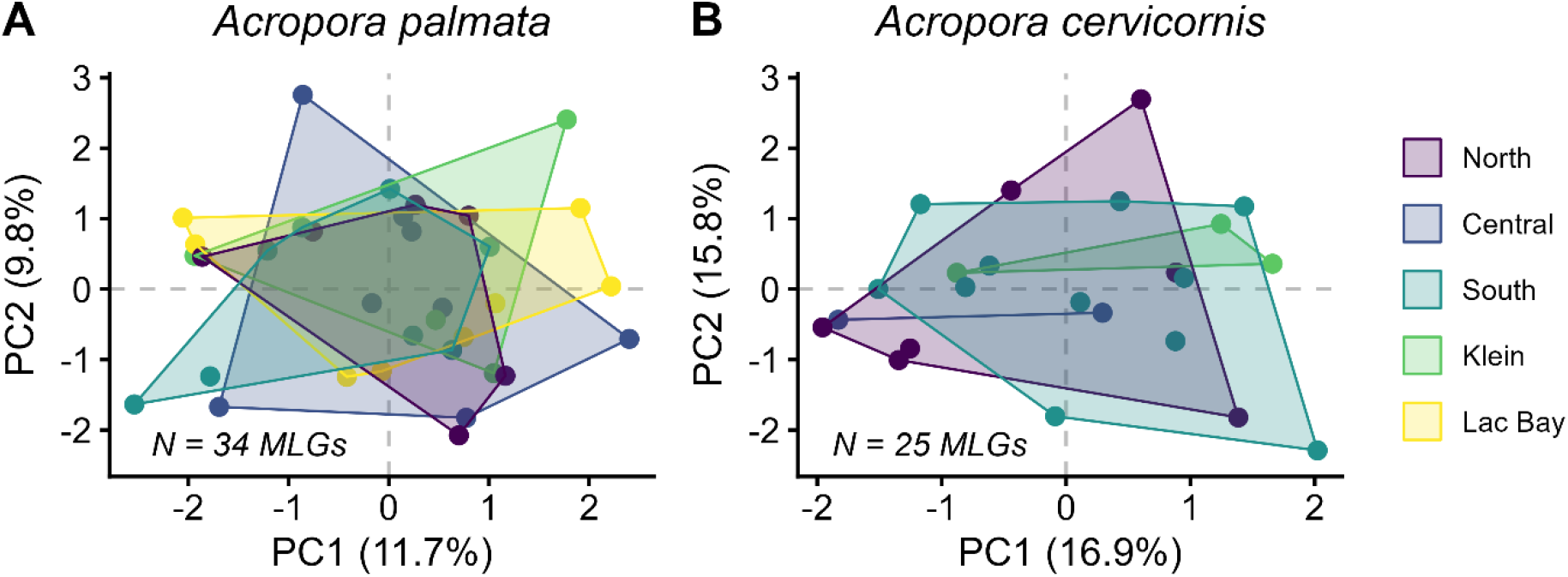
Principal Component Analysis (PCA) for *Acropora cervicornis* (A) and *Acropora palmata* (B), grouped by potential sub-populations around Bonaire. Each point represents a unique Multi-Locus Genotype (MLG). Eleven microsatellite loci informed the placement of each MLG in the PCA for *A. palmata,* and six loci informed MLG placement for *A. cervicornis*; rare cases of missing data were replaced using imputation. *A. cervicornis* data excluded Lac Bay because the only individual collected there (lineage G01) had a relatively high frequency of triploidy across loci. Sample sizes were small for *A. cervicornis* Central (n = 2) and Klein (n = 3), making the analyses of *A. cervicornis* low-powered and primarily exploratory.

**Table 3.**
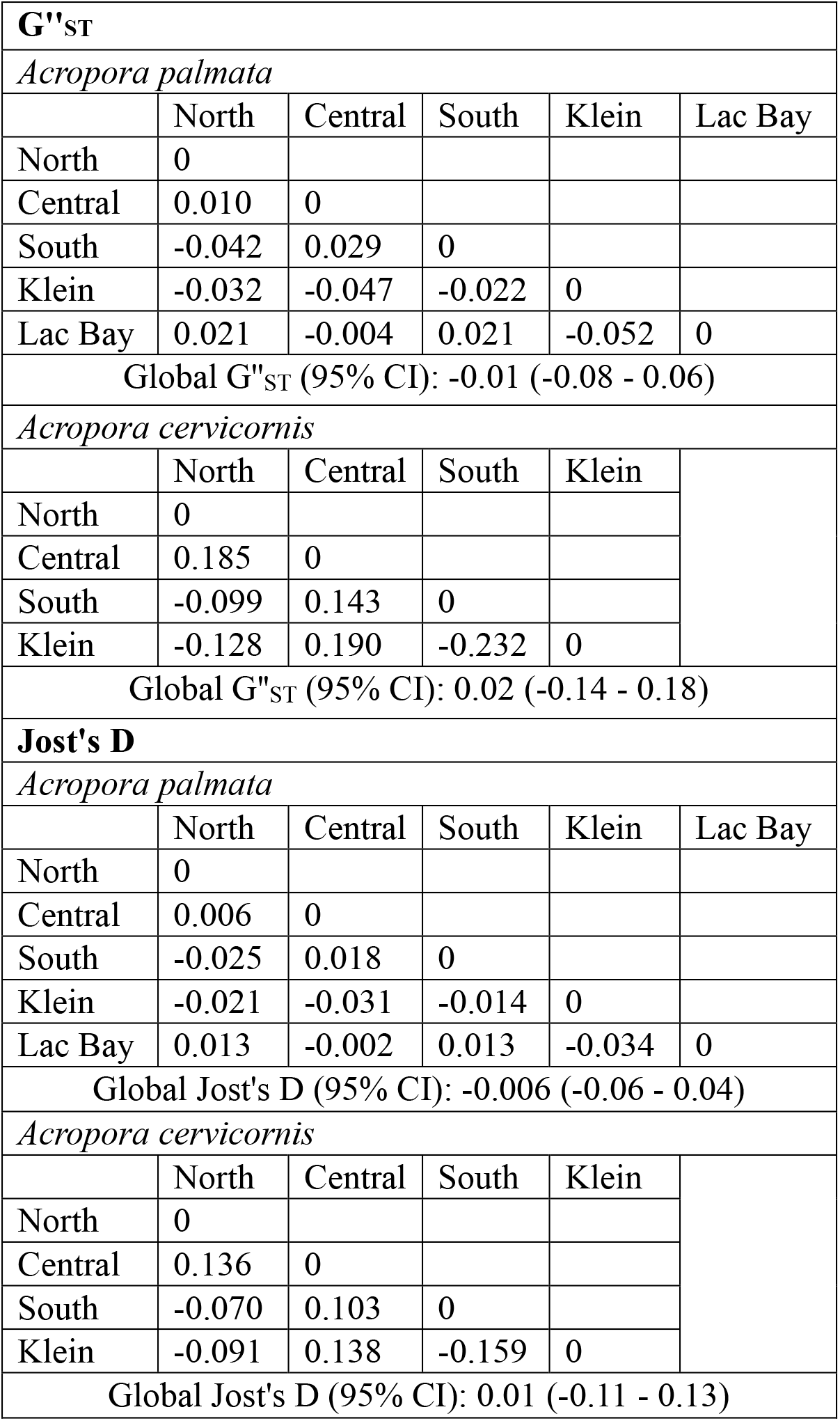
Pairwise and global comparisons of potential sub-populations in *Acropora palmata* and *Acropora cervicornis* for nearness to fixation (G’’_ST_) and allelic differentiation (Jost’s D). Global values comapre all potential subpopulations and were deemed not significantly greater than zero if 95% confidence interval (presented in parentheses) bounded zero. *A. cervicornis* analyses excluded Lac Bay because the only individual collected there (lineage G01) had a relatively high frequency of triploidy across loci. Sample sizes were small for *A. cervicornis* Central (n = 2) and Klein (n = 3), making the analyses of *A. cervicornis* low-powered and primarily exploratory.

| **G"<sub>ST</sub>** |  |  |  |  |  |
|---|---|---|---|---|---|
| *Acropora palmata* |  |  |  |  |  |
|  | North | Central | South | Klein | Lac Bay |
| North | 0 |  |  |  |  |
| Central | 0.010 | 0 |  |  |  |
| South | -0.042 | 0.029 | 0 |  |  |
| Klein | -0.032 | -0.047 | -0.022 | 0 |  |
| Lac Bay | 0.021 | -0.004 | 0.021 | -0.052 | 0 |
| Global G"<sub>ST</sub> (95% CI): -0.01 (-0.08 - 0.06) |  |  |  |  |  |
| *Acropora cervicornis* |  |  |  |  |  |
|  | North | Central | South | Klein |  |
| North | 0 |  |  |  |  |
| Central | 0.185 | 0 |  |  |  |
| South | -0.099 | 0.143 | 0 |  |  |
| Klein | -0.128 | 0.190 | -0.232 | 0 |  |
| Global G"<sub>ST</sub> (95% CI): 0.02 (-0.14 - 0.18) |  |  |  |  |  |
| **Jost's D** |  |  |  |  |  |
| *Acropora palmata* |  |  |  |  |  |
|  | North | Central | South | Klein | Lac Bay |
| North | 0 |  |  |  |  |
| Central | 0.006 | 0 |  |  |  |
| South | -0.025 | 0.018 | 0 |  |  |
| Klein | -0.021 | -0.031 | -0.014 | 0 |  |
| Lac Bay | 0.013 | -0.002 | 0.013 | -0.034 | 0 |
| Global Jost's D (95% CI): -0.006 (-0.06 - 0.04) |  |  |  |  |  |
| *Acropora cervicornis* |  |  |  |  |  |
|  | North | Central | South | Klein |  |
| North | 0 |  |  |  |  |
| Central | 0.136 | 0 |  |  |  |
| South | -0.070 | 0.103 | 0 |  |  |
| Klein | -0.091 | 0.138 | -0.159 | 0 |  |
| Global Jost's D (95% CI): 0.01 (-0.11 - 0.13) |  |  |  |  |  |

**Table 4.** Relatedness estimates (TrioML) and coefficients of inbreeding (F_IS_) and for each sub-population. *A. cervicornis* analyses excluded Lac Bay because the only individual collected there (lineage G01) had a relatively high frequency of triploidy across loci. Sample sizes were small for *A. cervicornis* Central (n = 2) and Klein (n = 3), making the analyses of *A. cervicornis* low-powered and primarily exploratory.

|  | **Relatedness** |  | **Inbreeding coefficient (F<sub>IS</sub>)** |  |
|---|---|---|---|---|
|  | *A. palmata* | *A. cervicornis* | *A. palmata* | *A. cervicornis* |
| North | 0.061 | 0.116 | -0.038 | 0.100 |
| Central | 0.081 | 0.298 | 0.037 | -0.056 |
| Klein | 0.027 | 0.101 | 0.139 | -0.037 |
| South | 0.053 | 0.098 | 0.168 | 0.013 |
| Lac Bay | 0.092 | NA | 0.106 | NA |
| Overall | 0.071 | 0.128 | 0.084 | 0.021 |

De novo exploration of potential structuring also indicated admixture, as there was no clear optimal number of clusters greater than one from either *find.clusters()* or Structure Selector methods (Supplemental Material S5, Fig. S2-S4). Specifically, in *find.clusters(),* the optimal number of clusters (i.e., lowest Bayesian Information Criterion (BIC) values) for both species was k = total MLG sample size, followed by k = 1 cluster for *A. palmata* and k = 2 clusters for *A. cervicornis*, though BIC values were low overall with little variation among k values (numbers of clusters). For Structure Selector, the optimal numbers of clusters (i.e., largest ΔK values) were k = 3 clusters for *A. palmata* and k = 10 clusters for *A. cervicornis*, however multiple other K values had relatively high ΔK well. We inspected STRUCTURE analysis outputs with K set equal to the optimal K identified in Structure Selector and found no differences in cluster membership among potential clusters or individuals. Thus, we concluded that there was no clear optimal number of clusters greater than one.

## Discussion

### Genetic diversity in gene bank

Reef Renewal Foundation Bonaire (RRFB) captured high genotypic richness (proportion of sampled individuals that are unique genotypes) in their gene bank, demonstrating the effectiveness of their gene banking strategy at maximizing the number of unique genotypes. In prior population genetics surveys of *Acropora spp.* throughout the Caribbean, a wide range in genotypic richness has been documented. For example, genotypic richness of *A. palmata* has ranged from 0.05 in Florida (Baums, Miller & Hellberg, 2006) to 1.0 in Saba (Japaud et al., 2019) and Navassa (Baums, Miller & Hellberg, 2006), while *A. cervicornis* ranged from 0.0125 in Guadeloupe (Japaud et al., 2015) to 1.0 in Yucatan, Mexico (Vollmer & Palumbi, 2007). Given the genotypic richness captured by RRFB, it is unlikely that Bonaire’s *Acropora spp.* populations are as clonal as those observed in Florida and Guadeloupe. In fact, one pair of *A. palmata* MLGs were collected from donor colonies only three meters apart from one another, highlighting that additional sampling even very close to already sampled colonies could yield new MLGs in Bonaire. To better understand the clonal structure of Bonaire’s *Acropora spp.* and therefore recommend a minimum distance between donor colonies for RRFB, future research could conduct standardized tissue collections (i.e., X individuals sampled within Y square meters) on patches of Bonaire’s wild (non-restored) *Acropora spp*.

RRFB’s gene banking captured a large percentage of Bonaire’s estimated asymptotic allelic diversity in *A. palmata* (88.9% of alleles captured with 34 MLGs) and *A. cervicornis* (74.7% of alleles captured with 25 MLGs) for the microsatellite loci we assessed. These results complement previous findings for coral and other animal taxa that sampling approximately 30 genetically unique individuals captures over 90% of a population’s microsatellite allelic diversity (Shearer, Porto & Zubillaga, 2009; Hale, Burg & Steeves, 2012). In comparison to previous studies on the genetic diversity of *Acropora spp.* in other locations across the Caribbean, the expected heterozygosity of *A. cervicornis* in RRFB’s gene bank was greater than nearly half of other populations, while *A. palmata* exhibited the lowest expected heterozygosity. It should be noted, though, that previous studies’ sampling designs aimed to estimate population-level genetic parameters, while RRFB aimed to maximize the number of unique genotypes. Thus, differences between the previous studies and the present one might represent differences in sampling design as well genetic diversity. Despite differences in sampling, both the expected heterozygosity and number of unique alleles of *A. palmata* in RRFB’s gene bank were similar to those of *A. palmata* from a single site (Tailor Made) in Bonaire in 2005 (Baums, Miller & Hellberg, 2005). This similarity in observed genetic diversity between one site 20 years ago and multiple sites in the present study supports the lack of strong genetic structuring (i.e., connectivity) that we observed among potential subpopulations. It also suggests that RRFB’s gene banking efforts successfully captured a comparable representation of the genetic diversity observed 20 years prior.

While RRFB captured a large percentage of Bonaire’s estimated *Acropora spp.* genetic diversity, we found that there were still many more *Acropora spp.* stands (i.e., individual colonies and dense thickets) that could be sampled to add potentially new genotypes to RRFB’s gene bank. More than 87% of Bonaire’s mapped stands, as of 2022, could still be potential new donor colonies for RRFB to continue enhancing its gene bank (i.e., 10.4% of mapped *A. palmata* stands and 12.5% of mapped *A. cervicornis* stands were RRFB collection sites). This is a conservative estimate, as it based off of mapping efforts that covered only 13% of the coastline and the unmapped sections likely have additional *Acropora spp.* stands.

### Minimal population structuring

We did not find evidence for genetic differences among potential subpopulations within Bonaire for either species, in terms of fixation (G’’_ST_, G_ST_, θ) or allelic differentiation (Jost’s D), indicating connectivity (i.e., admixture) around the island. While analyses for *A. cervicornis* were primarily exploratory due to low and unbalanced sample sizes in the potential subpopulations (e.g., Central = 2, Klein = 3), *A. cervicornis* exhibited similar patterns as *A. palmata,* which had greater statistical power. Both *A. palmata* and *A. cervicornis* exhibited relatedness estimates indicative of individuals being unrelated (<0.125), cousins (0.125), or half-siblings (0.25) within potential subpopulations and overall. While *A. palmata* exhibited greater inbreeding coefficients (F_IS_) than *A. cervicornis*, bootstrapping analyses indicated that observed relatedness values were not greater than expected for either species, meaning there was likely minimal inbreeding or non-random mating occurring. Further, both species exhibited inbreeding coefficients ≤0.17 within and among potential subpopulations, which is comparable to those documented in other scleractinian corals and marine invertebrates overall (Baums, 2008; Olsen et al., 2020). The lack of apparent inbreeding in Bonaire further points to connectivity around the island, as individuals do not appear to be restricted to mating with others within their subpopulations. Collectively, these results provide evidence of minimal population structuring (i.e., admixture) of Bonaire’s *A. palmata* and *A. cervicornis*. Thus, while the potential subpopulations were spatially separated, they were not genetically distinct from one another.

Our results support previous work that demonstrated high connectivity and diversity within populations of *Acropora spp.* in other locations (Vollmer & Palumbi, 2007; Porto-Hannes et al., 2015; Drury et al., 2016, 2017) and showed that Bonaire’s *Acropora spp.* are part of a larger interconnected eastern Caribbean population that spans multiple countries (Baums, Miller & Hellberg, 2005). Bonaire’s MPA managers, STINAPA, can therefore likely treat the island’s *Acropora spp.* as one genetically connected management unit for each species. To build off of our results, future research should explore more fine-scale genetic structuring and selection within Bonaire’s population by increasing sample sizes and utilizing high-throughput sequencing methods (Garcia Reyes & Schizas, 2010; Drury et al., 2017). Such work would either reinforce our results of connectivity or reveal weak population genetic structure, which could add nuance to the delineation of management units and alter RRFB’s breeding and outplanting program.

### Recommendations for restoration

Bonaire’s *Acropora spp.* have been strongly impacted by bleaching-induced mortality in conjunction with rising ocean temperatures in recent years (van Duyl, 1985; Sommer et al., 2011; Meesters et al., 2025), and RRFB’s gene banking efforts have likely saved some MLGs from fully disappearing. We recommend that RRFB continue collecting new lineages from wild colonies to preserve additional genetic diversity in their gene bank and maximize genetic diversity in their outplanting. Continued collection of new genotypes could increase the genetic diversity in RRFB’s restoration program by increasing allelic richness closer to the asymptotic maximum, increasing expected heterozygosity (particularly for *A. palmata*), and providing additional unique MLGs for sexual breeding. Across coral taxa, we recommend that coral restoration programs prioritize genotyping their lineages and maintain at least 30 MLGs to capture the majority of the population’s microsatellite allelic diversity, as has been found in *A. palmata, Montastrea cavernosa, Orbicella faveolata, Pocillopora damicornis,* and other animal taxa (Shearer, Porto & Zubillaga, 2009; Hale, Burg & Steeves, 2012).

For RRFB’s sexual breeding program, which is aimed at increasing genetic diversity and resilience, we recommend crossing pairs of MLGs in the gene bank that have low relatedness and avoiding pairs with relatedness ≥0.125 (i.e., cousins), such as the two *A. cervicornis* MLGs from Central Bonaire (relatedness = 0.298). By identifying unique MLGs, our results lay the groundwork for future research to assess variation among MLGs in resilience to stressors and heritability of fitness traits. We recommend that this research be conducted, so that RRFB might identify and prioritize MLGs for breeding that have greater fitness to create a more resilient population. As the restoration of other coral taxa becomes more common (Page, Muller & Vaughan, 2018; Bayraktarov et al., 2020), it will likewise be important to assess the genetic diversity and identify resilient genotypes, or lack thereof (Bell et al., 2026), of other species as well. The MLGs safeguarded by RRFB represent an important resource that could play a role in assisted gene flow as a restoration strategy for *Acropora spp.* throughout the Caribbean (Hagedorn et al., 2021; Baker et al., 2025) and serve as valuable contributions to a global biorepository network (Hagedorn et al., 2025).

We also suggest that an open-access database for archiving microsatellite genotyping data would be highly useful for coral restoration practice and research by improving the accessibility and reproducibility of microsatellite genotyping data and analyses. Species’ MLGs should not be directly compared between studies because allele identification can vary among laboratories based on PCR conditions and scoring criteria. However, a microsatellite database would be valuable for meta-analyses to calculate and compare descriptive statistics such as allelic richness and expected heterozygosity of coral taxa across regions, while controlling for sample size of individuals and microsatellite loci. While such a database does not exist, other pre-existing databases related to *Acropora spp.* restoration and genetics have been created, such as Coral Sample Registry (Moura et al., 2021), *A. cervicornis* Data Coordination Hub (Kiel et al., 2023), and STAGdb (Kitchen et al., 2020). A microsatellite database would complement these existing resources in cases when microsatellites are chosen because of their cost-effectiveness, as well as for archiving microsatellite data that has already been produced across multiple major restoration programs (Carne, Kaufman & Scavo, 2016; O’Donnell et al., 2018; Miller et al., 2019; Calle-Triviño et al., 2020).

## Conclusions

Here, we provided the most comprehensive assessment to date of the genetic diversity in Bonaire of two critically endangered coral species (Crabbe et al., 2022; Miller et al., 2022). We demonstrated that the coral restoration program led by Reef Renewal Foundation Bonaire (RRFB) captured high genotypic richness of *Acropora spp.*in their gene bank and relatively high allelic richness in comparison to the estimated asymptotic maximum. Bonaire’s *Acropora spp.* appeared to be well-connected around the island, as we did not detect genetic differentiation among spatially separated potential subpopulations and a lack of inbreeding and non-random mating. Collectively, our results demonstrated the effectiveness of RRFB’s lineage collection strategies to capture high genetic diversity and can be used to inform future initiatives on coral sexual breeding by RRFB and assisted gene flow beyond Bonaire.

## Acknowledgements

We thank Stichting Nationale Parken Bonaire (STINAPA) and the Bonaire and Dutch governments for permitting this research (Nature permit issued by Openbaar Lichaam Bonaire, archive number R&O2023000162). We thank the Convention on International Trade in Endangered Species (CITES) for permitting the collection and export of *Acropora spp.* samples (permit no. BQ/2022/002, BQ/2024/003), as well as the staff and volunteers at Reef Renewal Foundation Bonaire past and present that have contributed to coral lineage collection and nursery maintenance. We thank Yueling Hao, Steve Miller, Cathryn Ramon, and Maia Erbes for support conducting laboratory work and troubleshooting. We thank Yueling Hao, Steve Miller, Scott Miller, Beatriz Shobe, and Thomas Kemenes van Uden for assistance in data analysis.

## Notes

https://doi.org/10.5281/zenodo.23219412

### Competing Interest Statement

The authors have declared no competing interest.

## References

Aronson RB, Precht WF. 2001. White-band disease and the changing face of Caribbean coral reefs. Hydrobiologia 460:25–38. DOI: 10.1007/978-94-017-3284-0_2.

Baker AC, Baums IB, Davies SW, Grottoli AG, Kenkel CD, Kitchen SA, Kuffner IB, Matz MV, Miller MW, Muller EM, Parkinson JE, Prada C, Shantz AA, Hooidonk R van, Winters S. 2025. Proactive assisted gene flow for Caribbean corals in an era of rapid coral reef decline. Science 389:344–347. DOI: 10.1126/science.adx5842.

Baums IB. 2008. A restoration genetics guide for coral reef conservation. Molecular Ecology 17:2796–2811. DOI: 10.1111/j.1365-294X.2008.03787.x.

Baums IB, Chamberland VF, Locatelli NS, Conn T. 2022. Maximizing Genetic Diversity in Coral Restoration Projects. In: van Oppen MJH, Aranda Lastra M eds. Coral Reef Conservation and Restoration in the Omics Age. Cham: Springer International Publishing, 35–53. DOI: 10.1007/978-3-031-07055-6_3.

Baums IB, Devlin-Durante MK, Brown L, Pinzón JH. 2009. Nine novel, polymorphic microsatellite markers for the study of threatened Caribbean acroporid corals. Molecular Ecology Resources 9:1155–1158. DOI: 10.1111/j.1755-0998.2009.02581.x.

Baums IB, Hughes CR, Hellberg ME. 2005. Mendelian microsatellite loci for the Caribbean coral Acropora palmata. Marine Ecology Progress Series 288:115–127. DOI: 10.3354/meps288115.

Baums IB, Kitchen SA. 2020. Acropora DNA extraction with Qiagen DNeasy tissue kit. DOI: 10.17504/protocols.io.bgjqjumw.

Baums IB, Miller MW, Hellberg ME. 2005. Regionally isolated populations of an imperiled Caribbean coral, Acropora palmata. Molecular Ecology 14:1377–1390. DOI: 10.1111/j.1365-294X.2005.02489.x.

Baums IB, Miller MW, Hellberg ME. 2006. Geographic Variation in Clonal Structure in a Reef-Building Caribbean Coral, Acropora Palmata. Ecological Monographs 76:503–519. DOI: 10.1890/0012-9615(2006)076[0503:GVICSI]2.0.CO;2.

Bayraktarov E, Banaszak AT, Maya PM, Kleypas J, Arias-González JE, Blanco M, Calle-Triviño J, Charuvi N, Cortés-Useche C, Galván V, Salgado MAG, Gnecco M, Guendulain-García SD, Delgado EAH, Moraga JAM, Maya MF, Quiroz SM, Cervantes SM, Morikawa M, Nava G, Pizarro V, Sellares-Blasco RI, Ramos SES, Cubero TV, Villalpando MF, Frías-Torres S. 2020. Coral reef restoration efforts in Latin American countries and territories. PLOS ONE 15:e0228477. DOI: 10.1371/journal.pone.0228477.

Becking LE, Meesters HWG. 2014. Bonaire Deep Reef Expedition I.

Bell SL, Shilling EN, Pantoni GS, Eckert RJ, Sturm AB, Sharp WC, Muller EM, Gilliam DS, Lirman D, Bourque AS, Voss JD. 2026. Genetic lineages do not predict stony coral tissue loss disease resistance or survivorship among coral restoration outplants in Florida. Restoration Ecology 34:e70234. DOI: 10.1111/rec.70234.

Blacket MJ, Robin C, Good RT, Lee SF, Miller AD. 2012. Universal primers for fluorescent labelling of PCR fragments—an efficient and cost-effective approach to genotyping by fluorescence. Molecular Ecology Resources 12:456–463. DOI: 10.1111/j.1755-0998.2011.03104.x.

Boström-Einarsson L, Babcock RC, Bayraktarov E, Ceccarelli D, Cook N, Ferse SCA, Hancock B, Harrison P, Hein M, Shaver E, Smith A, Suggett D, Stewart-Sinclair PJ, Vardi T, McLeod IM. 2020. Coral restoration – A systematic review of current methods, successes, failures and future directions. PLOS ONE 15:e0226631. DOI: 10.1371/journal.pone.0226631.

Bruckner AW. 2003. Proceedings of the Caribbean Acropora Workshop: Potential Application of the U.S. Endangered Species Act as a Conservation Strategy. Miami, Florida: NOAA. URL: https://repository.library.noaa.gov/view/noaa/435

Calle-Triviño J, Rivera-Madrid R, León-Pech MG, Cortés-Useche C, Sellares-Blasco RI, Aguilar-Espinosa M, Arias-González JE. 2020. Assessing and genotyping threatened staghorn coral *Acropora cervicornis* nurseries during restoration in southeast Dominican Republic. PeerJ 8:e8863. DOI: 10.7717/peerj.8863.

Canty SWJ, Fox G, Rowntree JK, Preziosi RF. 2021. Genetic structure of a remnant Acropora cervicornis population. Scientific Reports 11:3523. DOI: 10.1038/s41598-021-83112-4.

Capel SLR, Bouzat JL, Catchen JM, Johnson JA, Dunn PO, Paige KN. 2022. Evaluating the genome-wide impacts of species translocations: the greater prairie-chicken as a case study. Conservation Genetics 23:179–191. DOI: 10.1007/s10592-021-01412-8.

Carne L, Kaufman L, Scavo K. 2016. Measuring success for Caribbean acroporid restoration: key results from ten years of work in southern Belize. In: Proceedings 13th International Coral Reef Symposium. Honolulu, HI, 17. URL: https://coralreefs.org/wp-content/uploads/2019/01/Session-42-B-2-Carne-ICRS-clean-and-final-ATB1.pdf

Chao A, Gotelli NJ, Hsieh TC, Sander EL, Ma KH, Colwell RK, Ellison AM. 2014. Rarefaction and extrapolation with Hill numbers: a framework for sampling and estimation in species diversity studies. Ecological Monographs 84:45–67. DOI: 10.1890/13-0133.1.

Chao A, Jost L, Chiang SC, Jiang Y-H, Chazdon RL. 2008. A Two-Stage Probabilistic Approach to Multiple-Community Similarity Indices. Biometrics 64:1178–1186. DOI: 10.1111/j.1541-0420.2008.01010.x.

Chessel D, Dufour AB, Thioulouse J. 2004. The ade4 package-I : One-table methods. *R News* 4. URL: https://journal.r-project.org/articles/RN-2004-002/RN-2004-002.pdf

Coker DJ, Wilson SK, Pratchett MS. 2014. Importance of live coral habitat for reef fishes. Reviews in Fish Biology and Fisheries 24:89–126. DOI: 10.1007/s11160-013-9319-5.

Crabbe J, Rodríguez-Martínez R, Villamizar E, Goergen L, Croquer A, Banaszak A. 2022. Acropora cervicornis. The IUCN Red List of Threatened Species 2022: e.T133381A165860142. *Available at* https://www.iucnredlist.org/es/species/133381/165860142#assessment-information (accessed May 6, 2025).

Cramer KL, Jackson JBC, Donovan MK, Greenstein BJ, Korpanty CA, Cook GM, Pandolfi JM. 2020. Widespread loss of Caribbean acroporid corals was underway before coral bleaching and disease outbreaks. Science Advances 6:eaax9395. DOI: 10.1126/sciadv.aax9395.

De Meyer K, MacRae DR, Raming J, Eckrich C, van der Ploeg J. 2022. Bonaire National Marine Park Management Plan 2022–2028.

Drury C, Dale KE, Panlilio JM, Miller SV, Lirman D, Larson EA, Bartels E, Crawford DL, Oleksiak MF. 2016. Genomic variation among populations of threatened coral: Acropora cervicornis. BMC Genomics 17:286. DOI: 10.1186/s12864-016-2583-8.

Drury C, Schopmeyer S, Goergen E, Bartels E, Nedimyer K, Johnson M, Maxwell K, Galvan V, Manfrino C, Lirman D. 2017. Genomic patterns in Acropora cervicornis show extensive population structure and variable genetic diversity. Ecology and Evolution 7:6188–6200. DOI: 10.1002/ece3.3184.

Dufresne F, Stift M, Vergilino R, Mable BK. 2014. Recent progress and challenges in population genetics of polyploid organisms: an overview of current state-of-the-art molecular and statistical tools. Molecular Ecology 23:40–69. DOI: 10.1111/mec.12581.

van Duyl F. 1985. Atlas of the living reefs of Curaçao and Bonaire (Netherlands Antilles). Ph.D. Thesis. Amsterdam: Vrije Universiteit. URL: https://linkinghub.elsevier.com/retrieve/pii/S0025326X24012748

Exposito-Alonso M, Booker TR, Czech L, Gillespie L, Hateley S, Kyriazis CC, Lang PLM, Leventhal L, Nogues-Bravo D, Pagowski V, Ruffley M, Spence JP, Toro Arana SE, Weiß CL, Zess E. 2022. Genetic diversity loss in the Anthropocene. Science 377:1431–1435. DOI: 10.1126/science.abn5642.

Gaither MR, Szabó Z, Crepeau MW, Bird CE, Toonen RJ. 2011. Preservation of corals in salt-saturated DMSO buffer is superior to ethanol for PCR experiments. Coral Reefs 30:329–333. DOI: 10.1007/s00338-010-0687-1.

Garcia Reyes J, Schizas NV. 2010. No two reefs are created equal: fine-scale population structure in the threatened coral species Acropora palmata and A. cervicornis. Aquatic Biology 10:69–83. DOI: 10.3354/ab00254.

García-Urueña R, Kitchen SA, Schizas NV. 2022. Fine scale population structure of Acropora palmata and Acropora cervicornis in the Colombian Caribbean. PeerJ 10:e13854. DOI: 10.7717/peerj.13854.

Goudet J, Jombart T. 2022. hierfstat: Estimation and Tests of Hierarchical F-Statistics. R package version 0.5–11, https://github.com/jgx65/hierfstat.

Hagedorn M, Page CA, O’Neil KL, Flores DM, Tichy L, Conn T, Chamberland VF, Lager C, Zuchowicz N, Lohr K, Blackburn H, Vardi T, Moore J, Moore T, Baums IB, Vermeij MJA, Marhaver KL. 2021. Assisted gene flow using cryopreserved sperm in critically endangered coral. Proceedings of the National Academy of Sciences 118:e2110559118. DOI: 10.1073/pnas.2110559118.

Hagedorn M, Zuchowicz N, Henley EM, Lager C, Perry R, Blackburn H, Bouwmeester J, Brunel O, Carter C, Rodriguez-Clark KM, Comizzoli P, Firchau B, Miller D, Moore J, Muller EM, O’Neil K, Quattrini AM, Zoccola D, Banaszak AT, Marhaver KL, Hobbs R, O’Brien JK, Daly J. 2025. Conservation of coral genetic diversity through a global biorepository network. BioScience:biaf117. DOI: 10.1093/biosci/biaf117.

Hale ML, Burg TM, Steeves TE. 2012. Sampling for Microsatellite-Based Population Genetic Studies: 25 to 30 Individuals per Population Is Enough to Accurately Estimate Allele Frequencies. PLOS ONE 7:e45170. DOI: 10.1371/journal.pone.0045170.

HDX, 2017.Bonaire, Sint Eustatius and Saba - Subnational Administrative Boundaries. Humanitarian Data Exchange, United Nations Office for the Coordination of Humanitarian Affairs (OCHA). URL: https://data.humdata.org/dataset/cod-ab-bes (accessed May 28, 2026).

Hedrick PW. 2005. A Standardized Genetic Differentiation Measure. Evolution 59:1633–1638. DOI: 10.1111/j.0014-3820.2005.tb01814.x.

Holl KD, Luong JC, Brancalion PHS. 2022. Overcoming biotic homogenization in ecological restoration. Trends in Ecology & Evolution 37:777–788. DOI: 10.1016/j.tree.2022.05.002.

Hsieh TC, Ma KH, Chao A. 2016. iNEXT: an R package for rarefaction and extrapolation of species diversity (Hill numbers). Methods in Ecology and Evolution 7:1451–1456. DOI: 10.1111/2041-210X.12613.

Jackson J, Donovan M, Cramer K, Lam W. 2014. Status and Trends of Caribbean Coral Reefs: 1970-2012. *Gland, Switzerland: Global Coral Reef Monitoring Network*, IUCN.

Japaud A, Bouchon C, Magalon H, Fauvelot C. 2019. Geographic distances and ocean currents influence Caribbean Acropora palmata population connectivity in the Lesser Antilles. Conservation Genetics 20:447–466. DOI: 10.1007/s10592-019-01145-9.

Japaud A, Bouchon C, Manceau J-L, Fauvelot C. 2015. High clonality in Acropora palmata and Acropora cervicornis populations of Guadeloupe, French Lesser Antilles. Marine and Freshwater Research 66:847–851. DOI: 10.1071/MF14181.

Johnson WE, Onorato DP, Roelke ME, Land ED, Cunningham M, Belden RC, McBride R, Jansen D, Lotz M, Shindle D, Howard J, Wildt DE, Penfold LM, Hostetler JA, Oli MK, O’Brien SJ. 2010. Genetic Restoration of the Florida Panther. Science 329:1641–1645. DOI: 10.1126/science.1192891.

Jombart T. 2008. adegenet: a R package for the multivariate analysis of genetic markers. Bioinformatics 24:1403–1405. DOI: 10.1093/bioinformatics/btn129.

Jombart T, Devillard S, Balloux F. 2010. Discriminant analysis of principal components: a new method for the analysis of genetically structured populations. BMC Genetics 11:94. DOI: 10.1186/1471-2156-11-94.

Jost L. 2008. *G*_ST_ and its relatives do not measure differentiation. Molecular Ecology 17:4015–4026. DOI: 10.1111/j.1365-294X.2008.03887.x.

Kemenes van Uden T, Meesters HWG, Dogruer G. 2023. Methods for Assessing Acropora spp. Populations in the Caribbean. URL: https://research.wur.nl/en/publications/methods-for-assessing-acropora-spp-populations-in-the-caribbean

Kiel PM, Formel N, Jankulak M, Baker AC, Cunning R, Gilliam DS, Kenkel CD, Langdon C, Lirman D, Lustic C, Maxwell K, Moulding AL, Moura A, Muller EM, Schopmeyer S, Winters RS, Enochs IC. 2023. Acropora Cervicornis Data Coordination Hub, an Open Access Database for Evaluating Genet Performance. Bulletin of Marine Science 99:119–136. DOI: 10.5343/bms.2022.0064.

Kitchen SA, Von Kuster G, Kuntz KLV, Reich HG, Miller W, Griffin S, Fogarty ND, Baums IB. 2020. STAGdb: a 30K SNP genotyping array and Science Gateway for Acropora corals and their dinoflagellate symbionts. Scientific Reports 10:12488. DOI: 10.1038/s41598-020-69101-z.

Lemopoulos A, Prokkola JM, Uusi-Heikkilä S, Vasemägi A, Huusko A, Hyvärinen P, Koljonen M-L, Koskiniemi J, Vainikka A. 2019. Comparing RADseq and microsatellites for estimating genetic diversity and relatedness — Implications for brown trout conservation. Ecology and Evolution 9:2106–2120. DOI: 10.1002/ece3.4905.

Li Y-L, Liu J-X. 2018. StructureSelector: A web-based software to select and visualize the optimal number of clusters using multiple methods. Molecular Ecology Resources 18:176–177. DOI: 10.1111/1755-0998.12719.

Manzello DP, Cunning R, Karp RF, Baker AC, Bartels E, Bonhag R, Borreil A, Bourque A, Brown KT, Bruckner AW, Corbett B, D’Alessandro M, Dahlgren C, Dilworth J, Geiger E, Gilliam DS, Gomez M, Hanson G, Harrell C, Hesley D, Huebner LK, Kenkel CD, Koch HR, Kuehl J, Kuffner IB, Ladd MC, Lee S, Lesneski KC, Lewan A, Lirman D, Liu G, Matsuda SB, Montoya-Maya PH, Moore J, Muller EM, Nedimyer K, Parkinson JE, Ruzicka R, Spadaro J, Spady BL, Stein J, Unsworth JD, Walter C, Wen ADE, Williams DE, Williams SD, Williamson OM. 2025. Heat-driven functional extinction of Caribbean Acropora corals from Florida’s Coral Reef. Science 390:361–366. DOI: 10.1126/science.adx7825.

Meesters EH, Van Der Geest M, Kemenes Van Uden T, Boman E, Butler E, Hylkema A, Lehwald M, Wulf K, Eckrich C, Francisca R. 2025. Conservation state of coral reefs and communities of the Caribbean Netherlands. Den Helder: Wageningen Marine Research. DOI: 10.18174/694539.

Meirmans PG. 2015. Seven common mistakes in population genetics and how to avoid them. Molecular Ecology 24:3223–3231. DOI: 10.1111/mec.13243.

Meirmans PG, Hedrick PW. 2011. Assessing population structure: FST and related measures. Molecular Ecology Resources 11:5–18. DOI: 10.1111/j.1755-0998.2010.02927.x.

Meirmans PG, Liu S, van Tienderen PH. 2018. The Analysis of Polyploid Genetic Data. Journal of Heredity 109:283–296. DOI: 10.1093/jhered/esy006.

Miller MW, Colburn PJ, Pontes E, Williams DE, Bright AJ, Serrano XM, Peters EC. 2019. Genotypic variation in disease susceptibility among cultured stocks of elkhorn and staghorn corals. PeerJ 7:e6751. DOI: 10.7717/peerj.6751.

Miller M, Crabbe J, Villamizar E, Croquer A, Banaszak A. 2022. Acropora palmata. IUCN Red List of Threatened Species 2022: e.T133006A165672016. *Available at* https://www.iucnredlist.org/en (accessed December 3, 2025).

Moura A, Beck B, Duffey R, McEachron L, Miller M, Moore J, Moulding A, Winters RS. 2021. Integrating Coral Restoration Data With a Novel Coral Sample Registry. Frontiers in Marine Science 8:700172. DOI: 10.3389/fmars.2021.700172.

Nei M. 1973. Analysis of Gene Diversity in Subdivided Populations. Proceedings of the National Academy of Sciences 70:3321–3323. DOI: 10.1073/pnas.70.12.3321.

Nylander-Asplin HF, Hill RL, Doerr JC, Greer L, Fogarty ND. 2021. Population dynamics and genotypic richness of threatened Acropora species and their hybrid in the U.S. Virgin Islands. Coral Reefs 40:965–971. DOI: 10.1007/s00338-021-02093-y.

O’Brien D, Aavik T, Fedorca A, Fischer MC, Goffaux R, Hoban S, Hollingsworth P, Hvilsom C, Jehle R, Stroil BK, Kershaw F, Klinga P, Kopatz A, Leigh DM, Paz-Vinas I, Robuchon M, Segelbacher G, Takacs V, Vernesi C, Laikre L. 2025. Restoring genetic diversity to facilitate the implementation of the EU Nature Restoration Law. Biological Conservation 303:110995. DOI: 10.1016/j.biocon.2025.110995.

O’Donnell KE, Lohr KE, Bartels E, Baums IB, Patterson JT. 2018. Acropora cervicornis genet performance and symbiont identity throughout the restoration process. Coral Reefs 37:1109–1118. DOI: 10.1007/s00338-018-01743-y.

Olsen KC, Ryan WH, Winn AA, Kosman ET, Moscoso JA, Krueger-Hadfield SA, Burgess SC, Carlon DB, Grosberg RK, Kalisz S, Levitan DR. 2020. Inbreeding shapes the evolution of marine invertebrates. Evolution 74:871–882. DOI: 10.1111/evo.13951.

van Oosterhout C, Hutchinson WF, Wills DPM, Shipley P. 2004. micro-checker: software for identifying and correcting genotyping errors in microsatellite data. Molecular Ecology Notes 4:535–538. DOI: 10.1111/j.1471-8286.2004.00684.x.

Page CA, Muller EM, Vaughan DE. 2018. Microfragmenting for the successful restoration of slow growing massive corals. Ecological Engineering 123:86–94. DOI: 10.1016/j.ecoleng.2018.08.017.

Pew J, Muir PH, Wang J, Frasier TR. 2015. related: an R package for analysing pairwise relatedness from codominant molecular markers. Molecular Ecology Resources 15:557–561. DOI: 10.1111/1755-0998.12323.

Porto-Hannes I, Zubillaga AL, Shearer TL, Bastidas C, Salazar C, Coffroth MA, Szmant AM. 2015. Population structure of the corals Orbicella faveolata and Acropora palmata in the Mesoamerican Barrier Reef System with comparisons over Caribbean basin-wide spatial scale. Marine Biology 162:81–98. DOI: 10.1007/s00227-014-2560-1.

Rolloos EF, Dogruer G, Meesters HWG. 2023. The distribution of Acropora spp. along the Leeward coast of Bonaire. Wageningen Marine Research. URL: https://library.wur.nl/WebQuery/wurpubs/628458

RRFB. 2024. Reef Renewal Bonaire 2024 Annual Report. URL: https://www.reefrenewalbonaire.org/about-us/annual-reports/

Shearer TL, Porto I, Zubillaga AL. 2009. Restoration of coral populations in light of genetic diversity estimates. Coral reefs (Online) 28:727–733. DOI: 10.1007/s00338-009-0520-x.

Sommer B, Harrison PL, Brooks L, Scheffers SR. 2011. Coral Community Decline at Bonaire, Southern Caribbean. Bulletin of Marine Science 87:541–565. DOI: 10.5343/bms.2010.1046.

Sunde J, Yıldırım Y, Tibblin P, Forsman A. 2020. Comparing the Performance of Microsatellites and RADseq in Population Genetic Studies: Analysis of Data for Pike (Esox lucius) and a Synthesis of Previous Studies. Frontiers in Genetics 11. DOI: 10.3389/fgene.2020.00218.

Taylor HR. 2015. The use and abuse of genetic marker-based estimates of relatedness and inbreeding. Ecology and Evolution 5:3140–3150. DOI: 10.1002/ece3.1541.

Vardi T, Hoot WC, Levy J, Shaver E, Winters RS, Banaszak AT, Baums IB, Chamberland VF, Cook N, Gulko D, Hein MY, Kaufman L, Loewe M, Lundgren P, Lustic C, MacGowan P, Matz MV, McGonigle M, McLeod I, Moore J, Moore T, Pivard S, Pollock FJ, Rinkevich B, Suggett DJ, Suleiman S, Viehman TS, Villalobos T, Weis VM, Wolke C, Montoya-Maya PH. 2021. Six priorities to advance the science and practice of coral reef restoration worldwide. Restoration Ecology 29:e13498. DOI: 10.1111/rec.13498.

Vollmer SV, Palumbi SR. 2007. Restricted Gene Flow in the Caribbean Staghorn Coral Acropora cervicornis: Implications for the Recovery of Endangered Reefs. Journal of Heredity 98:40–50. DOI: 10.1093/jhered/esl057.

Wang J. 2007. Triadic IBD coefficients and applications to estimating pairwise relatedness. Genetics Research 89:135–153. DOI: 10.1017/S0016672307008798.

Wei X, Xu Y, Lyu L, Xiao Z, Wang S, Yang T, Jiang M. 2023. Impacts of ecological restoration on the genetic diversity of plant species: A global meta-analysis. Journal of Applied Ecology 60:1149–1160. DOI: 10.1111/1365-2664.14390.

Weir BS, Cockerham CC. 1984. Estimating F-Statistics for the Analysis of Population Structure. Evolution 38:1358–1370. DOI: 10.2307/2408641.

Williams DE, Miller MW, Baums IB. 2014. Cryptic changes in the genetic structure of a highly clonal coral population and the relationship with ecological performance. Coral Reefs 33:595–606. DOI: 10.1007/s00338-014-1157-y.

Williams D, Nedimyer K, Bright A, Ladd M. 2024. Genotypic inventory and impact of the 2023 marine heatwave on Acropora palmata (elkhorn coral) populations in the Upper Florida Keys, USA: 2020–2023. URL: https://repository.library.noaa.gov/view/noaa/60881

Winter DJ. 2012. mmod: an R library for the calculation of population differentiation statistics. Molecular Ecology Resources 12:1158–1160. DOI: 10.1111/j.1755-0998.2012.03174.x.

Young C, Schopmeyer S, Lirman D. 2012. A Review of Reef Restoration and Coral Propagation Using the Threatened Genus Acropora in the Caribbean and Western Atlantic. Bulletin of Marine Science 88:1075–1098. DOI: 10.5343/bms.2011.1143.

Zimmerman SJ, Aldridge CL, Oyler-McCance SJ. 2020. An empirical comparison of population genetic analyses using microsatellite and SNP data for a species of conservation concern. BMC Genomics 21:382. DOI: 10.1186/s12864-020-06783-9.

