## Supplemental Material for "Genetic diversity of endangered *Acropora spp.* corals in Bonaire: implications for restoration and gene banking"

Supplemental Material S1-S10

**Supplemental Material S1. Comparison of DNA Shield and salt-saturated DMSO as preservatives for *Acropora cervicornis***

**Table S1. Comparison of DNA yields (µg/mL) from five *A. cervicornis* samples that had replicates separately preserved in DNA Shield and salt-saturated DMSO.** DNA concentrations of zero indicate when the values were too low to be read by the Qubit (e.g., "Out of Range").

| Lineage | Nursery tree code | Preservative | DNA concentration (µg/mL) |
|---|---|---|---|
| G03 | BN | DMSO | 3.58 |
| G04 | BN | DMSO | 0.00 |
| G05 | BN | DMSO | 0.00 |
| G09 | BN | DMSO | 23.95 |
| G09 | GB | DMSO | 31.90 |
| G03 | BN | DNA Shield | 31.72 |
| G04 | BN | DNA Shield | 20.72 |
| G05 | BN | DNA Shield | 14.45 |
| G09 | BN | DNA Shield | 22.37 |
| G09 | GB | DNA Shield | 5.99 |

### Supplemental Material S2. PCR primer sequences

**Table S2. PCR primer sequences for microsatellite loci and universal tails used in multiplex PCR.** Each multiplex PCR included microsatellite forward primers (with universal primer sequence at start), microsatellite reverse primers, and fluorescently tagged universal primers. Microsatellite locus primer sequences were developed and published by Baums, Hughes & Hellberg (2005) and Baums et al. (2009), and universal primer sequences were developed and published by Blacket et al. (2012).

| Locus ID | Universal primer | Direction | Sequence 5' to 3' |
|---|---|---|---|
| **Mix A** | | | |
| 182 | Tail A | Forward | GCCTCCCTCGCGCCATCCCACAACTCACACTCTGC |
| | | Reverse | ACGCGGAAATAGTGATGCTC |
| 0585 | Tail C | Forward | CAGGACCAGGCTACCGTGTGGGCAAAGTTGATTTCCTG |
| | | Reverse | TGAGACCCAAAGAAGTGGC |
| 1195 | Tail D | Forward | CGGAGAGCCGAGAGGTGCACCCGACGATTCGAAAGC |
| | | Reverse | TGACAAGGGATCAAATCAAGTTC |
| **Mix B** | | | |
| 5047 | Tail A | Forward | GCCTCCCTCGCGCCATCGGGTATATTGCGAGTACG |
| | | Reverse | GCAGGATCGTTCATTTGTATGC |
| 207 | Tail B | Forward | GCCTTGCCAGCCCGCATCCACGCCCAAACAATGTA |
| | | Reverse | CTATTCGCTACCCACGCTTC |
| 181 | Tail C | Forward | CAGGACCAGGCTACCGTGTTCTCCACATGCAAACAAACA |
| | | Reverse | GCCAGGATAGCGGATAATGA |
| 6212 | Tail D | Forward | CGGAGAGCCGAGAGGTGTTACAAGGTGCATAGGACCC |
| | | Reverse | CCTCTCCATCAGCCATTTCC |
| **Mix C** | | | |
| 0513 | Tail A | Forward | GCCTCCCTCGCGCCAAGCAAGACAGGAATGTGGG |
| | | Reverse | CCGGAAGCAACCTCATTGG |
| 2637 | Tail B | Forward | GCCTTGCCAGCCCGCGCGGCGATAGTAATTGGTC |
| | | Reverse | TGGTGGTTCAGCTTTGGC |
| 1490 | Tail C | Forward | CAGGACCAGGCTACCGTGTTCCTCACTTCGGTTGGGC |
| | | Reverse | ACGTTCCTTCATACCAAGAGC |
| 166 | Tail D | Forward | CGGAGAGCCGAGAGGTGTCTACCCGCAATTTTCATCA |
| | | Reverse | CGCTCTCCTATGTTCGATTG |
| **Fluorescently labeled universal primers** | | | |
| - | Tail A | - | FAM-GCCTCCCTCGCGCCA |
| - | Tail B | - | VIC-GCCTTGCCAGCCCGC |
| - | Tail C | - | NED-CAGGACCAGGCTACCGTG |
| - | Tail D | - | PET-CGGAGAGCCGAGAGGTG |

Baums IB, Hughes CR, Hellberg ME. 2005. Mendelian microsatellite loci for the Caribbean coral Acropora palmata. Marine Ecology Progress Series 288:115–127. DOI: 10.3354/meps288115.

Baums IB, Devlin-Durante MK, Brown L, Pinzón JH. 2009. Nine novel, polymorphic microsatellite markers for the study of threatened Caribbean acroporid corals. Molecular Ecology Resources 9:1155–1158. DOI: 10.1111/j.1755-0998.2009.02581.x.

Blacket MJ, Robin C, Good RT, Lee SF, Miller AD. 2012. Universal primers for fluorescent labelling of PCR fragments—an efficient and cost-effective approach to genotyping by fluorescence. *Molecular Ecology Resources* 12:456–463. DOI: 10.1111/j.1755-0998.2011.03104.x.

**Supplemental Material S3. Explanation of methods for assessing confidence in Multi-Locus Genotype identification (MLG).**

We assessed confidence in MLG identity using three methods in the poppr package (Kamvar et al. 2014). First, we determined that unique Multi-Locus Genotypes (MLGs) could be identified from four loci in *A. palmata* and five loci in *A. cervicornis,* via genotype accumulation curves (Fig. S1). In practice, we assessed 11 loci in *A. palmata* and six loci in *A. cervicornis.*

Second, we estimated the probability of obtaining identical MLGs in different lineages due to sexual reproduction or chance, as opposed to asexual cloning, using the $P_{sex}$ statistic and loci in Hardy Weinber Equilibrium (HWE, an assumption of the analysis). For *A. palmata* MLGs (nine loci in HWE), $P_{sex}$ was less than $4^{-5}$. For *A. cervicornis* (six loci in HWE), $P_{sex}$ was less than $2^{-3}$ for all MLGs except one. For the *A. cervicornis* lineage G01, $P_{sex}$ was 0.40, though this lineage only had available diploid data at four loci while loci 181, 1490, and 6212 had triploid data that could not be included in $P_{sex}$ analyses. Due to the triploid nature of *A. cervicornis* lineage G01, we determined this lineage to be genetically distinct from all others. Overall, it was unlikely that any lineages would have shared the same MLG identity via sexual reproduction or chance, as opposed to asexual cloning.

Third, we assessed if there was low genetic distance (Bruvo's distance threshold < 0.1) between any of the MLGs that would be indicative of somatic mutations within a single MLG, as has been observed in other populations of *A. palmata* and *A. cervicornis* (Porto-Hannes et al. 2015, Japaud et al. 2015, Canty et al. 2021). No *A. palmata* MLGs appeared to differ due to somatic mutations, with all MLGs differing in at least six loci. No *A. cervicornis* MLGs appeared to differ due to somatic mutations, with all MLGs differing in at least three alleles.

References

Canty SWJ, Fox G, Rowntree JK, Preziosi RF (2021) Genetic structure of a remnant Acropora cervicornis population. Sci Rep 11:3523.

Japaud A, Bouchon C, Manceau J-L, Fauvelot C (2015) High clonality in Acropora palmata and Acropora cervicornis populations of Guadeloupe, French Lesser Antilles. Mar Freshw Res 66:847–851.

Kamvar ZN, Tabima JF, Grünwald NJ (2014) Poppr: an R package for genetic analysis of populations with clonal, partially clonal, and/or sexual reproduction. PeerJ 2:e281.

Porto-Hannes I, Zubillaga AL, Shearer TL, Bastidas C, Salazar C, Coffroth MA, Szmant AM (2015) Population structure of the corals Orbicella faveolata and Acropora palmata in the Mesoamerican Barrier Reef System with comparisons over Caribbean basin-wide spatial scale. Mar Biol 162:81–98.

**Figure S1. Genotype accumulation curves depicting number of Multi-Locus Genotypes identified as a function of number of microsatellite loci used, for *Acropora palmata* (A) and *A. cervicornis* (B).**

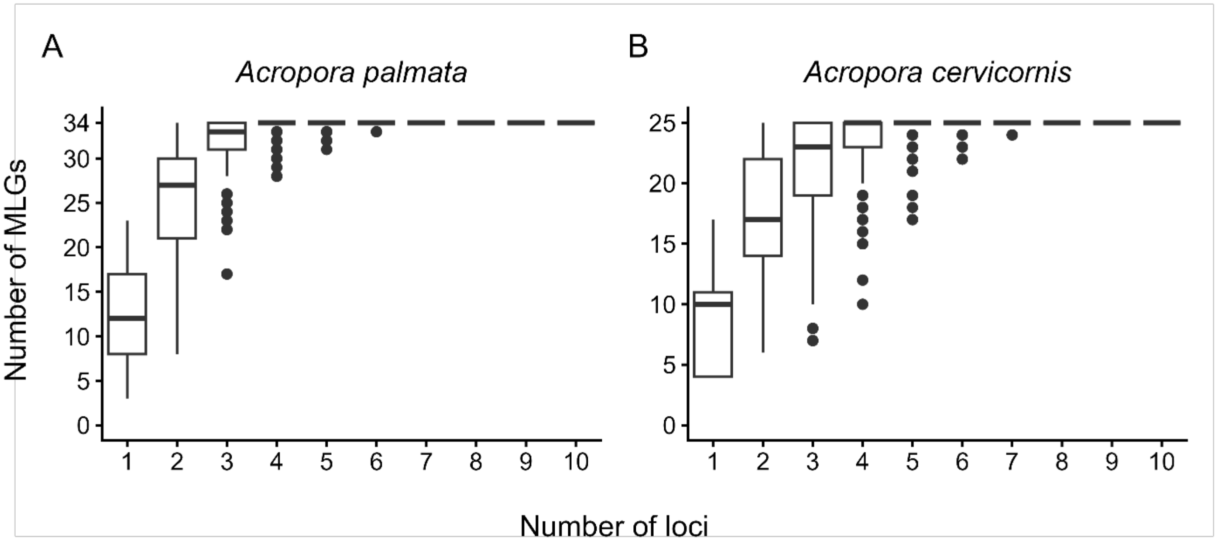

**Supplemental Material S4. Explanation of literature search methods for comparing reported genetic diversity metrics of other *Acropora spp.* populations.**

To search the literature for population genetic studies on *A. palmata* and *A. cervicornis* that reported genetic diversity metrics based on microsatellites, we searched the Web of Science Core Collection database on 02 March, 2026, using the Advanced Search option with the query TS = ("Acropora cervicornis" OR "Acropora palmata" AND "microsatellite" OR "allel*" OR "heterozygosity" OR "population" OR "structure" OR "genetic*" OR "diversit*"). We filtered for peer-reviewed journal publications from 2005 to present, because 2005 was the year the first Caribbean *Acropora spp.* microsatellites were published (Baums et al. 2005). All resulting publications were in English. We screened the first 100 returned publications for reported results on number of unique microsatellite alleles and expected heterozygosity.

Baums IB, Hughes CR, Hellberg ME (2005) Mendelian microsatellite loci for the Caribbean coral Acropora palmata. Mar Ecol Prog Ser 288:115–127. https://doi.org/10.3354/meps288115

**Supplemental Material S5. Interpretation of de novo exploration of clustering among Bonaire's *Acropora palmata* and *Acropora cervicornis* using *find.cluster()* and Structure Selector.**

To interpret the optimal number of clusters using *find.cluster()* in adegenet (Jombart 2008, Jombart et al. 2010), we generated and inspected Bayesian Information Criterion (BIC) plots. To generate these plots, we set the maximum number of clusters equal to the sample size of total MLGs, to assess all possible numbers of clusters. We set the number of PCs retained equal to 50, to ensure that all possible PCs were included (Jombart & Collins 2022). Typical BIC plots have a convex curve, in which the number of clusters with the lowest BIC value represents the optimal number of clusters for the dataset (Jombart et al. 2010). Alternative to typical BIC plots, we observed concave curves for both coral species (Fig. S2), with lowest BIC values at the maximum (N Multi-Locus Genotypes) number of clusters, which is not biologically relevant. The next lowest BIC value for *A. palmata* was at k=1 cluster. For *A. cervicornis,* k = 2 had the next-lowest BIC value, though k=1-5 all had approximately the same BIC value. Overall, our BIC values were relatively low, with little variation among numbers of clusters, relative to those in Jombart et al (2010). Thus, the BIC plots generated in *find.clusters()* did not provide strong support for any biologically relevant optimal number of clusters greater than one.

To assess the optimal number of clusters using Structure Selector (Li & Liu 2018), we generated and inspected plots of ΔK (Evanno et al. 2005), in which the largest ΔK value indicates the optimal number of clusters (K). To inform Structure Selector, we ran STRUCTURE v.2.3.4 (Pritchard et al. 2000) analyses with loci in Hardy Weinberg Equilibrium (HWE, *A. palmata*: nine loci, *A. cervicornis*: six loci), because HWE is an assumption of this analysis. We tested the number of clusters (K) ranging from one to the total number of MLGs, with 20 replicates per K. Each replicate run had a 50,000 burn-in period length, plus 100,000 iterations after burn-in.

The output ΔK plots from Structure Selector (Fig. S3) indicated that *A. palmata* optimal K = 3 (i.e., mean 11.3 MLGs per cluster) and *A. cervicornis* optimal K = 10 (i.e., mean 2.3 MLGs per cluster), however multiple other K values had relatively high as ΔK well. Next, we inspected STRUCTURE analysis outputs when K was set equal to the optimal K identified in ΔK plots (Fig. S4). Overall, the STRUCTURE analysis outputs did not show any differences in cluster membership among potential clusters or individuals, when set to the optimal K.

In summary, both *find.clusters()* (Jombart 2008, Jombart et al. 2010) and Structure Selector (Li & Liu 2018) de novo methods indicated admixture, as there was no clear optimal number of clusters greater than one.

References

Evanno G, Regnaut S, Goudet J (2005) Detecting the number of clusters of individuals using the software structure: a simulation study. Mol Ecol 14:2611–2620.

Jombart T (2008) Adegenet: a R package for the multivariate analysis of genetic markers. Bioinformatics 24:1403–1405.

Jombart T, Collins C (2022) A tutorial for Discriminant Analysis of Principal Components (DAPC) using adegenet 2.1.6

Jombart T, Devillard S, Balloux F (2010) Discriminant analysis of principal components: a new method for the analysis of genetically structured populations. BMC Genet 11:94.

Li Y-L, Liu J-X (2018) StructureSelector: A web-based software to select and visualize the optimal number of clusters using multiple methods. Mol Ecol Resour 18:176–177.

**Figure S2. Bayesian Information Criterion (BIC) plots for de novo exploration of clustering, via *find.clusters()*.**

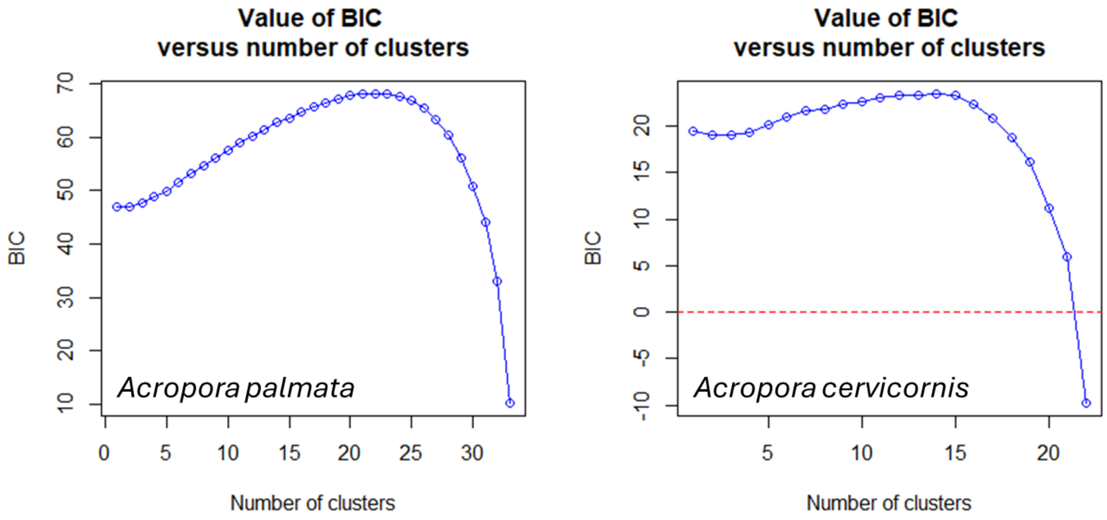


**Figure S3. ΔK plots for de novo exploration of clustering, via Structure Selector.**

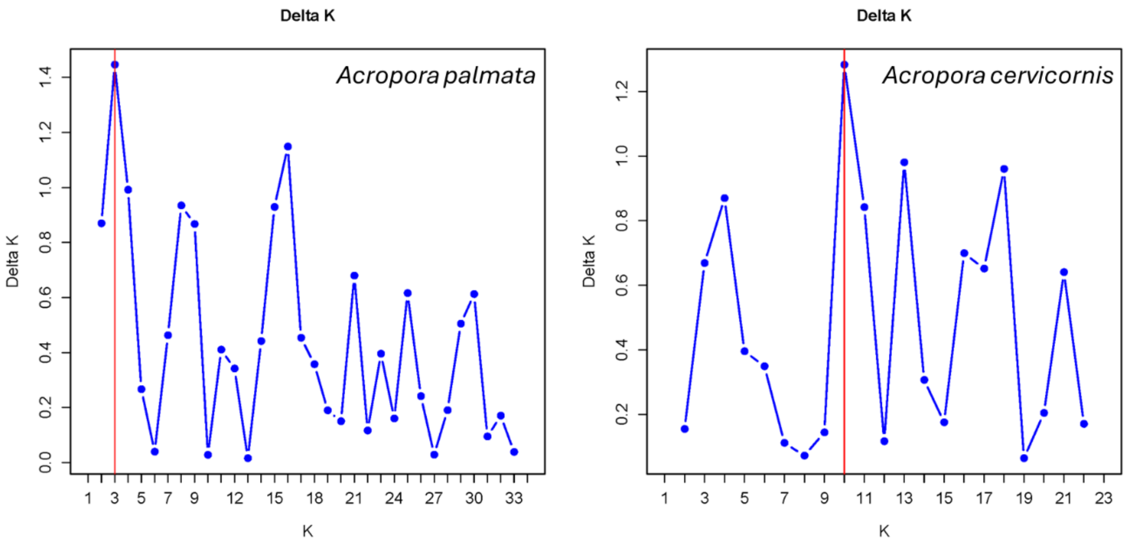

**Figure S4. STRUCTURE analysis outputs when K was set equal to the optimal K identified in ΔK plots (*Acropora palmata*: K = 3, *A. cervicornis*: K = 10).** Potential sub-populations are North (1), Central (2), South (3), Klein (4), Lac Bay (5). *A. cervicornis* analyses excluded Lac Bay because the only individual collected there (lineage G01) had a relatively high frequency of triploidy across loci. Sample sizes were small for *A. cervicornis* Central (n = 2) and Klein (n = 3), making the analyses of *A. cervicornis* low-powered and primarily exploratory.

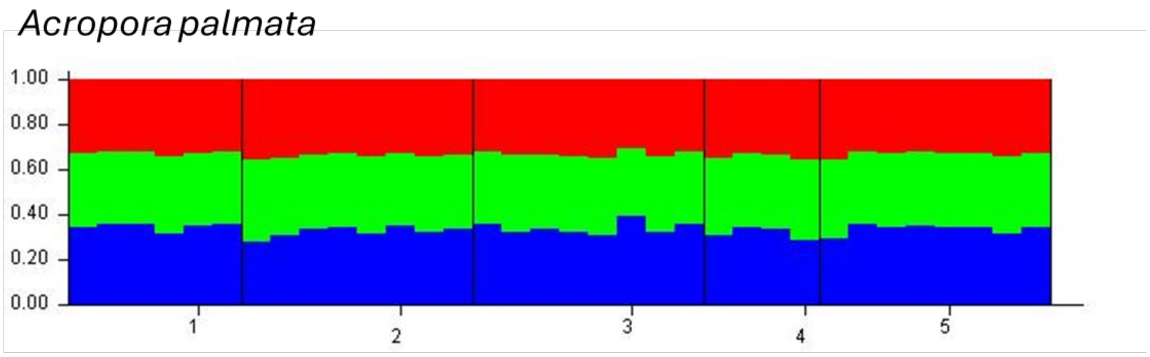


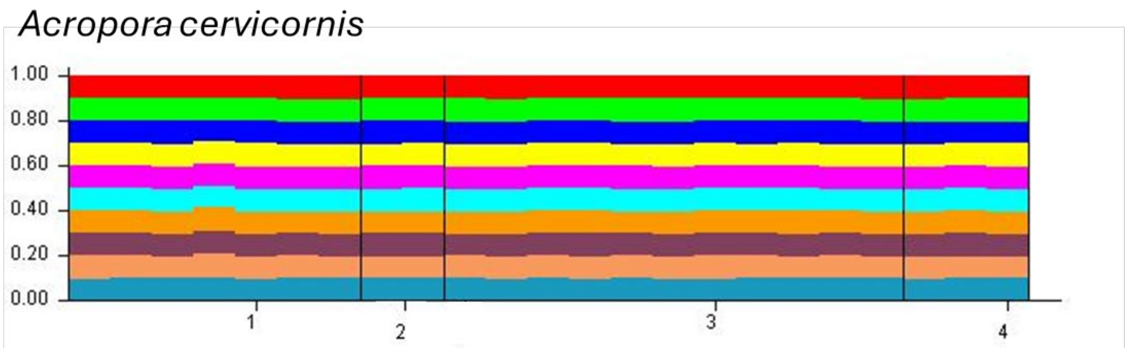

**Supplemental Material S6. Explanation of microsatellite locus screening for data quality control.**

Loci were successfully genotyped across ≥85% of lineages (i.e., <15% missing data) in all 11 loci in *A. palmata* (182, 0585, 1195, 5047, 207, 181, 6212, 0513, 2637, 1490, 166) and in six loci in *A. cervicornis* (182, 0585, 5047, 181, 1490, 166). For data quality control, we screened each microsatellite locus with three steps. First, we visually re-inspected peak calls across all 11 loci in both species in Geneious Prime. Second, we checked for linkage disequilibrium across loci with <15% missing data using poppr (Agapow & Burt 2001, Kamvar et al. 2014). There was no apparent linkage disequilibrium among loci for *A. palmata* ($\bar{r}_d$ = 0.006, p = 0.270) or *A. cervicornis* ($\bar{r}_d$ = 0.034, p = 0.049).

Third, we assessed for possible null alleles in loci with <15% missing data by testing for significant deviations from Hardy-Weinberg Equilibrium (HWE) with pegas (Paradis 2010) and by estimating null allele frequencies with PopGenReport (Adamack & Gruber 2014). Loci 207 and 166 in *A. palmata* significantly deviated from HWE (Table 1), however null allele frequencies were not significantly greater than zero (lower bounds of 95% confidence intervals <0) in either species. To further inspect loci 207 and 166 in *A. palmata* for null alleles, we estimated null allele frequencies in ML-Null Freq (Kalinowski & Taper 2006) and Micro-checker (van Oosterhout et al. 2004) as well. Both programs estimated null allele frequencies consistent with PopGenReport, though Micro-checker suggested that locus 166 might have null alleles. We tested if the results of this study were altered if locus 166 was excluded from *A. palmata* and found no difference. Therefore, we retained loci 207 and 166 in *A. palmata* for subsequent analyses. Locus 1195 was flagged in its original publication as having relatively high estimates of null alleles (Baums et al. 2009), however we found no evidence of null alleles and included 1195 in subsequent analyses.

References

Adamack AT, Gruber B (2014) PopGenReport: simplifying basic population genetic analyses in R. Methods Ecol Evol 5:384–387.

Agapow P-M, Burt A (2001) Indices of multilocus linkage disequilibrium. Mol Ecol Notes 1:101–102.

Baums IB, Devlin-Durante MK, Brown L, Pinzón JH (2009) Nine novel, polymorphic microsatellite markers for the study of threatened Caribbean acroporid corals. Mol Ecol Resour 9:1155–1158.

Kalinowski ST, Taper ML (2006) Maximum likelihood estimation of the frequency of null alleles at microsatellite loci. Conserv Genet 7:991–995.

Kamvar ZN, Tabima JF, Grünwald NJ (2014) Poppr: an R package for genetic analysis of populations with clonal, partially clonal, and/or sexual reproduction. PeerJ 2:e281.

van Oosterhout C, Hutchinson WF, Wills DPM, Shipley P (2004) Micro-checker: software for identifying and correcting genotyping errors in microsatellite data. Mol Ecol Notes 4:535–538.

Supplemental Material S1-S10

Paradis E (2010) Pegas: an R package for population genetics with an integrated–modular approach. Bioinformatics 26:419–420.

**Supplemental Material S7. Locus rarefactions of number of unique alleles as a function of Multi-locus Genotype sample size, across all evaluated loci per species.**

**Figure S5. *Acropora palmata* rarefactions (11 microsatellite loci evaluated).**

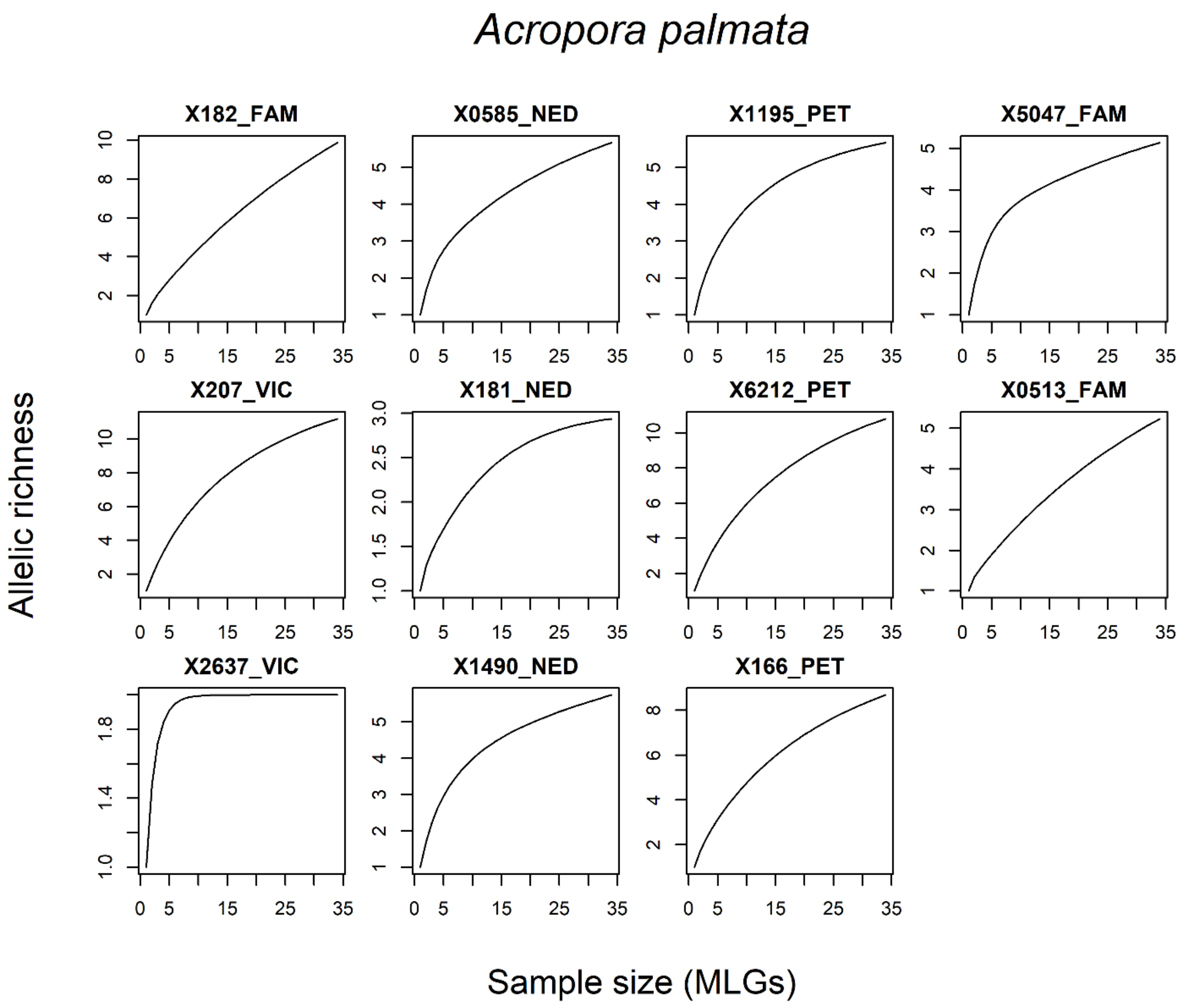

**Figure S6.** ***Acropora cervicornis*** **rarefactions (six microsatellite loci evaluated).**

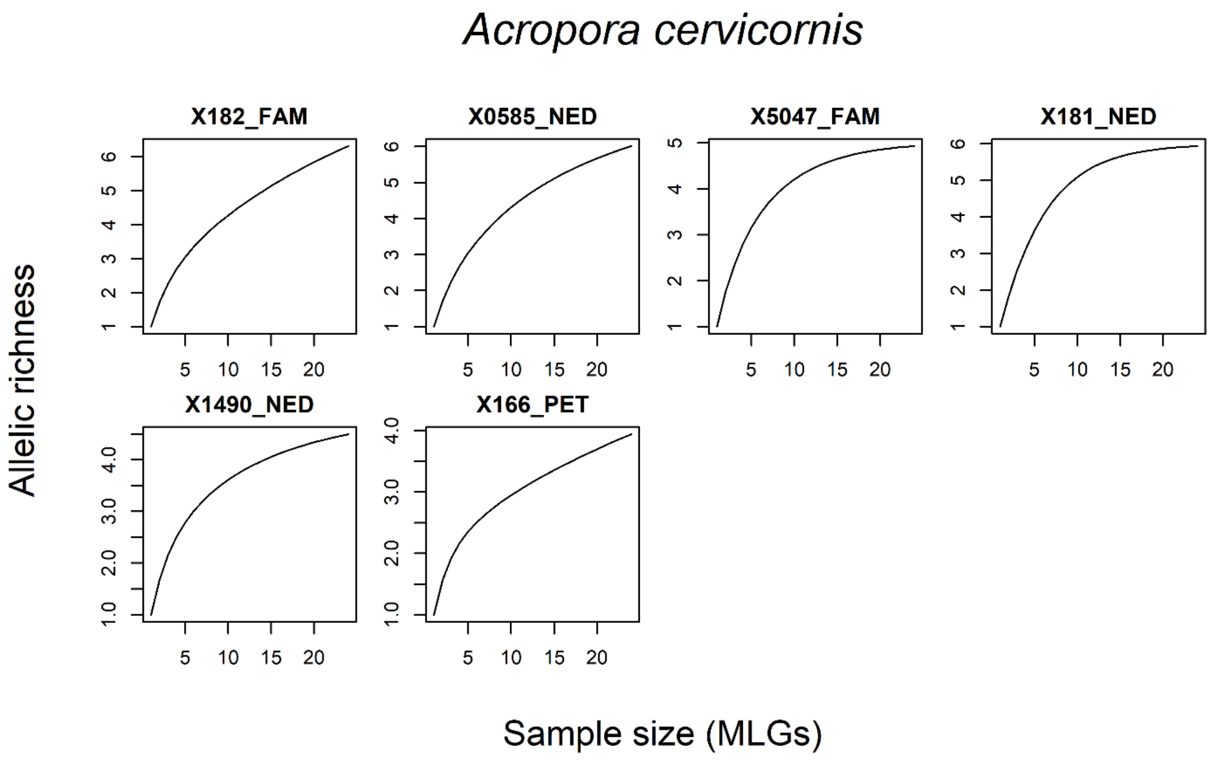

**Supplemental Material S8. Comparison of *Acropora spp.* donor-colony locations to mapping surveys of *Acropora spp.* stands.**

**Figure S7. Maps of locations of donor colonies where Reef Renewal Foundation Bonaire collected lineages.** Points are color-coded by whether the collection location was from a stand (i.e., individual colony or dense thicket) of *Acropora spp.* mapped in 2022 surveys by Wageningen Marine Research and STINAPA (Kemenes van Uden et al. 2023, Rolloos et al. 2023) that surveyed 146 hectares total shelf area (approximately 13% coastline). Orange represents stands identified in mapping surveys; blue represents non-mapped stands. Map shapefile sourced from Humanitarian Data Exchange (HDX, 2017).

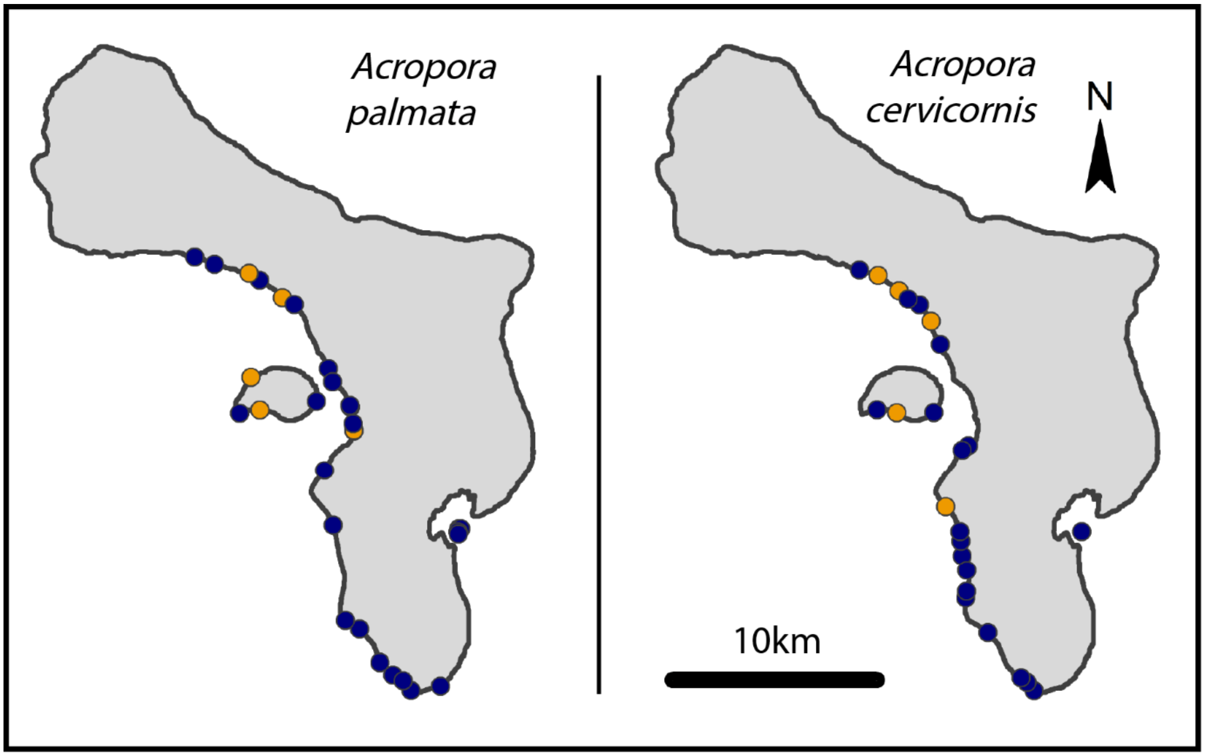


References

HDX. 2017.Bonaire, Sint Eustatius and Saba - Subnational Administrative Boundaries. Humanitarian Data Exchange, United Nations Office for the Coordination of Humanitarian Affairs (OCHA). URL: https://data.humdata.org/dataset/cod-ab-bes (accessed May 28, 2026).

Kemenes van Uden T, Meesters HWG, Dogruer G (2023) Methods for Assessing Acropora spp. Populations in the Caribbean.

Rolloos EF, Dogruer G, Meesters HWG (2023) The distribution of Acropora spp. along the Leeward coast of Bonaire. Wageningen Marine Research.

### Supplemental Material S9. Pairwise comparisons of $G_{ST}$ and θ

**Table S3. Pairwise comparisons of nearness to fixation among potential sub-populations in *Acropora palmata* and *A. cervicornis,* using $G_{ST}$ (Nei 1973) and θ (Weir and Cockerham 1984).**

**$G_{ST}$**

*Acropora palmata*

| | North | Central | South | Klein | Lac Bay |
|---|---|---|---|---|---|
| North | 0 | | | | |
| Central | 0.002 | 0 | | | |
| South | -0.008 | 0.006 | 0 | | |
| Klein | -0.006 | -0.008 | -0.004 | 0 | |
| Lac Bay | 0.004 | -0.001 | 0.004 | -0.009 | 0 |

*Acropora cervicornis*

| | North | Central | South | Klein |
|---|---|---|---|---|
| North | 0 | | | |
| Central | 0.029 | 0 | | |
| South | -0.013 | 0.023 | 0 | |
| Klein | -0.017 | 0.031 | -0.031 | 0 |

**θ**

*Acropora palmata*

| | North | Central | South | Klein | Lac Bay |
|---|---|---|---|---|---|
| North | 0 | | | | |
| Central | 0.002 | 0 | | | |
| South | -0.022 | 0.005 | 0 | | |
| Klein | -0.018 | -0.023 | -0.021 | 0 | |
| Lac Bay | 0.004 | -0.006 | 0.000 | -0.029 | 0 |

*Acropora cervicornis*

| | North | Central | South | Klein |
|---|---|---|---|---|
| North | 0 | | | |
| Central | 0.035 | 0 | | |
| South | -0.030 | 0.036 | 0 | |
| Klein | -0.044 | 0.067 | -0.064 | 0 |

**Supplemental Material S10. Histograms of bootstrapped expected relatedness values (TrioML) within and across potential sub-populations.** Red line indicates observed relatedness.

**Figure S8.** ***Acropora palmata*** **relatedness histograms.**

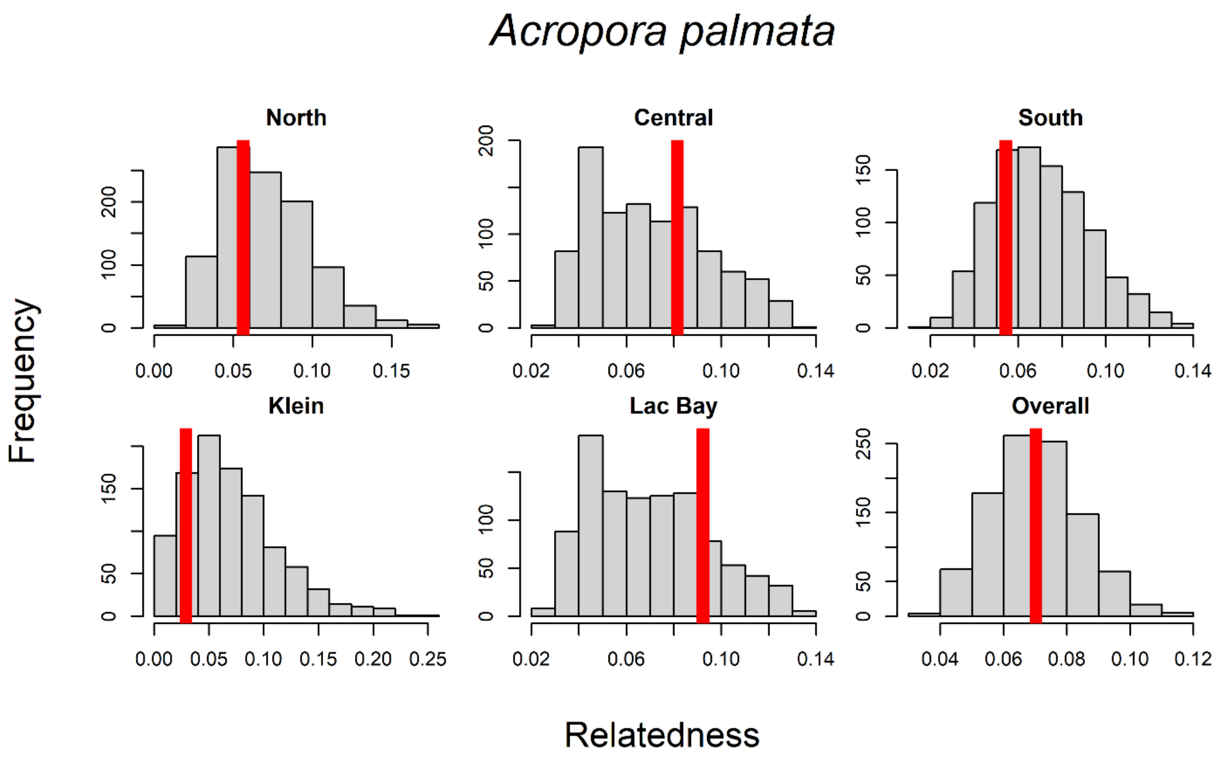

**Figure S9. *Acropora cervicornis* relatedness histograms.**

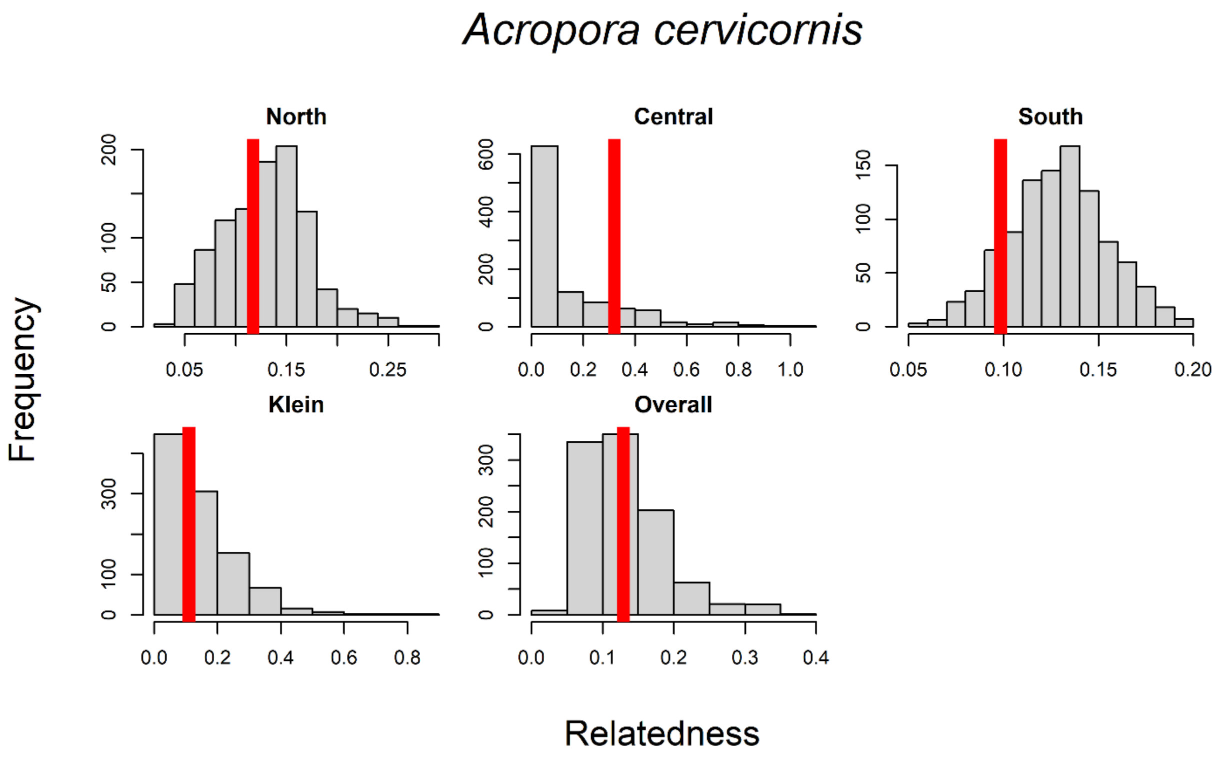
